# Phosphorylation barcodes direct biased chemokine signaling at CXCR3

**DOI:** 10.1101/2023.03.14.532634

**Authors:** Dylan S. Eiger, Jeffrey S. Smith, Tujin Shi, Tomasz Maciej Stepniewski, Chia-Feng Tsai, Christopher Honeycutt, Noelia Boldizsar, Julia Gardner, Carrie D. Nicora, Ahmed M. Moghieb, Kouki Kawakami, Issac Choi, Kevin Zheng, Anmol Warman, Priya Alagesan, Nicole M. Knape, Ouwen Huang, Justin D. Silverman, Richard D. Smith, Asuka Inoue, Jana Selent, Jon M. Jacobs, Sudarshan Rajagopal

**Affiliations:** Department of Biochemistry, Duke University, Durham, NC, 27710, USA; Department of Dermatology, Massachusetts General Hospital, Boston, MA, 02114, USA; Department of Dermatology, Brigham and Women’s Hospital, Boston, MA, 02115, USA; Department of Dermatology, Beth Israel Deaconess Medical Center, Boston, MA, 02215, USA; Dermatology Program, Boston Children’s Hospital, Boston, MA, 02115, USA; Harvard Medical School, Boston, MA, 02115, USA; Biological Sciences Division, Pacific Northwest National Laboratory, Richland, WA, 99354, USA; Research Programme on Biomedical Informatics (GRIB), Department of Experimental and Health Sciences of Pompeu Fabra University (UPF)-Hospital del Mar Medical Research Institute (IMIM), Barcelona, 08003, Spain; Trinity College, Duke University, Durham, NC, 27710, USA; R&D Research, GlaxoSmithKline, Collegeville, PA, 19426, USA; Department of Pharmaceutical Sciences, Tohoku University, Sendai, 980-8577, Japan; Department of Medicine, Duke University, Durham, NC 27710 USA; Department of Biomedical Engineering, Duke University, Durham, NC, 27710, USA; College of Information Sciences and Technology, The Pennsylvania State University, University Park, PA, 16802, USA

**Author notes:** = these authors contributed equally (Dylan S. Eiger, Jeffrey S. Smith). = corresponding author (Sudarshan Rajagopal). = lead contact (Sudarshan Rajagopal).

**Keywords:** beta-arrestin, G protein-coupled receptor, biased agonism, chemokine, CXCR3, phosphoproteomics, chemotaxis, MAP kinase, mass spectrometry, molecular dynamics

## Abstract

G protein-coupled receptor (GPCR) biased agonism, the activation of some signaling pathways over others, is thought to largely be due to differential receptor phosphorylation, or “phosphorylation barcodes.” At chemokine receptors, ligands act as “biased agonists” with complex signaling profiles, which contributes to the limited success in pharmacologically targeting these receptors. Here, mass spectrometry-based global phosphoproteomics revealed that CXCR3 chemokines generate different phosphorylation barcodes associated with differential transducer activation. Chemokine stimulation resulted in distinct changes throughout the kinome in global phosphoproteomic studies. Mutation of CXCR3 phosphosites altered β-arrestin conformation in cellular assays and was confirmed by molecular dynamics simulations. T cells expressing phosphorylation-deficient CXCR3 mutants resulted in agonist- and receptor-specific chemotactic profiles. Our results demonstrate that CXCR3 chemokines are non-redundant and act as biased agonists through differential encoding of phosphorylation barcodes and lead to distinct physiological processes.

## INTRODUCTION

G protein-coupled receptors (GPCRs) are the most common transmembrane receptors in the human genome and the target of approximately one third of all FDA-approved drugs (Hauser et al., 2017). GPCRs elicit cellular responses by coupling to heterotrimeric G proteins, recruiting GPCR kinases (GRKs), and binding to β- arrestin adaptor proteins (Smith and Rajagopal, 2016). Certain GPCR ligands can promote or inhibit different GPCR conformational states, leading to distinct G protein or β-arrestin signaling outputs; i.e. display “biased agonism”. Efforts are underway to design biased agonists that preferentially activate certain signaling pathways to maximize clinical efficacy and reduce off-target effects (Pupo et al., 2016). However, the molecular determinants of biased signaling and the degree to which different ligands can modulate intracellular signaling cascades remain unclear.

Altering intracellular GPCR amino acid residue phosphorylation patterns can lead to different signaling events and is one mechanism for encoding biased agonism (Busillo et al., 2010; Butcher et al., 2011; Dwivedi-Agnihotri et al., 2020; Latorraca et al., 2020; Nobles et al., 2011). For example, preventing phosphorylation of certain residues impairs receptor endocytosis but not β-arrestin recruitment (Oakley et al., 1999). Specific GPCR phosphorylation patterns also differentially alter the affinity of GPCR-β-arrestin interactions (Bouzo-Lorenzo et al., 2016; Jung et al., 2017; Lee et al., 2016; Mayer et al., 2019; Sente et al., 2018). GPCR agonists are thought to regulate β-arrestin function by encoding distinct phosphorylation events through selective interaction with different GRKs (Busillo et al., 2010; Butcher et al., 2011; Inagaki et al., 2015; Komolov and Benovic, 2018; Nobles et al., 2011). This “phosphorylation barcode hypothesis” is supported by mutagenesis studies in both cellular and animal models (Bradley et al., 2020; Kaya et al., 2020; Kliewer et al., 2019; Mann et al., 2020; Marsango et al., 2022; Scarpa et al., 2021; Zhou et al., 2017). Biophysical data also support that different C-terminal phosphorylation patterns induce distinct β-arrestin conformational states (Dwivedi-Agnihotri *et al*., 2020; Lee *et al*., 2016; Mayer *et al*., 2019; Nobles *et al*., 2011; Nuber et al., 2016; Yang et al., 2015; Yang et al., 2017), and may expose β-arrestin binding sites for some downstream effectors but not others (Latorraca et al., 2020). However, it is unclear if different C-terminal residues are phosphorylated or if the same residues are phosphorylated at differing stoichiometric ratios, a distinction critical to understanding how GPCR is mechanistically encoded. In addition, few studies have identified distinct phosphopeptides or associated changes in phosphorylation barcodes with changes in physiology (Busillo et al., 2010; Butcher et al., 2011; Nobles et al., 2011). There remains limited evidence that specific phosphopeptide patterns promote changes in receptor signaling with downstream physiological effects, and understanding how ligands generate such signaling profiles is critical to understanding cellular signal transduction.

The physiological relevance of endogenous biased signaling can be difficult to assess as the majority of GPCR biased agonists are synthetic. However, many endogenous biased agonists have been identified in the chemokine system (Corbisier et al., 2015; Rajagopal et al., 2013), which consists of approximately 20 receptors and 50 chemokine ligands (Eiger et al., 2021; Griffith et al., 2014; Kufareva et al., 2015). Unlike other GPCR subfamilies, chemokine receptors are promiscuous and often bind multiple chemokines with high affinity (Allen et al., 2007; Zlotnik and Yoshie, 2012). For example, the chemokine receptor CXCR3, primarily expressed on activated T cells, binds three endogenous ligands, CXCL9, CXCL10, and CXCL11, and plays an important role in inflammatory diseases and cancer (Chow et al., 2019; Kuo et al., 2018; Smith et al., 2018c). Like most other chemokine receptors, CXCR3 signals through both Gαi family G proteins and β-arrestins (Colvin et al., 2006; Colvin et al., 2004; Smith et al., 2017). CXCL11 is β-arrestin-biased compared to CXCL9 and CXCL10, with each chemokine displaying distinct profiles of G protein signaling, β-arrestin recruitment and receptor internalization (Rajagopal *et al*., 2013; Zheng et al., 2022). Synthetic CXCR3 biased agonists have shown distinct physiological effects in a mouse model of allergic contact dermatitis, with a β-arrestin-biased agonist promoting inflammation through increased T cell recruitment (Smith et al., 2018a). CXCR3 is well-suited for studying the mechanisms underlying biased agonism and its physiological impact.

It is unclear how receptors with multiple endogenous ligands encode divergent cellular signaling and function. Here we demonstrate that endogenous chemokines of CXCR3 encode unique phosphorylation barcode ensembles (different phosphopeptides at different stoichiometries). These differential phosphorylation ensembles lead to different patterns of transducer and kinome activation with subsequent distinctive chemotactic patterns. Through mutagenic studies, we determined that CXCR3 biased signaling is encoded through the receptor core and differential phosphorylation of the receptor C-terminal tail.

## RESULTS

### CXCR3 chemokines promote different receptor phosphorylation barcode ensembles

CXCR3 chemokines (CXCL9, CXCL10, CXCL11) promote different levels of receptor phosphorylation (Colvin et al., 2004; Smith et al., 2017). However, it is not known whether this is due to differences of phosphorylation levels at the same sites or at different sites on the C-terminus, or both. To determine if the endogenous CXCR3 chemokines produce different phosphorylation barcode ensembles (different site patterns and levels of those patterns), we utilized state-of-the-art mass spectrometry with combinatorial phosphopeptide reference libraries with heavy isotope labeled reference standards corresponding to potential CXCR3 serine and threonine phosphorylation patterns as previously described (Tsai et al., 2019). This approach allowed us to not only identify but also quantify the relative abundance of specific phosphopeptides after chemokine stimulation. Wild-type human CXCR3-overexpressing HEK293 cells were stimulated with CXCL9, CXCL10, CXCL11 or vehicle control, followed by tryptic digestion and tandem mass tag (TMT) labelling, allowing samples to be pooled and greatly improving measurement precision as well as eliminating variability from batch effects **(Figure 1A and 1B)**. After ion metal affinity chromatography (IMAC) enrichment, TMT-labeled peptides were analyzed using liquid chromatography and tandem mass spectrometry (LC-MS/MS) for phosphopeptide identification (**Figure 1B**). Phosphosites of interest were further validated by targeted proteomics with the addition of a synthetic library of 128 heavy isotope-labeled C-terminal phosphopeptides prior to IMAC enrichment. This enabled us to confidently differentiate and quantify adjacent phosphosites, providing high-resolution insights into the ensemble of receptor phosphopeptides following chemokine stimulation.

**Figure 1:**
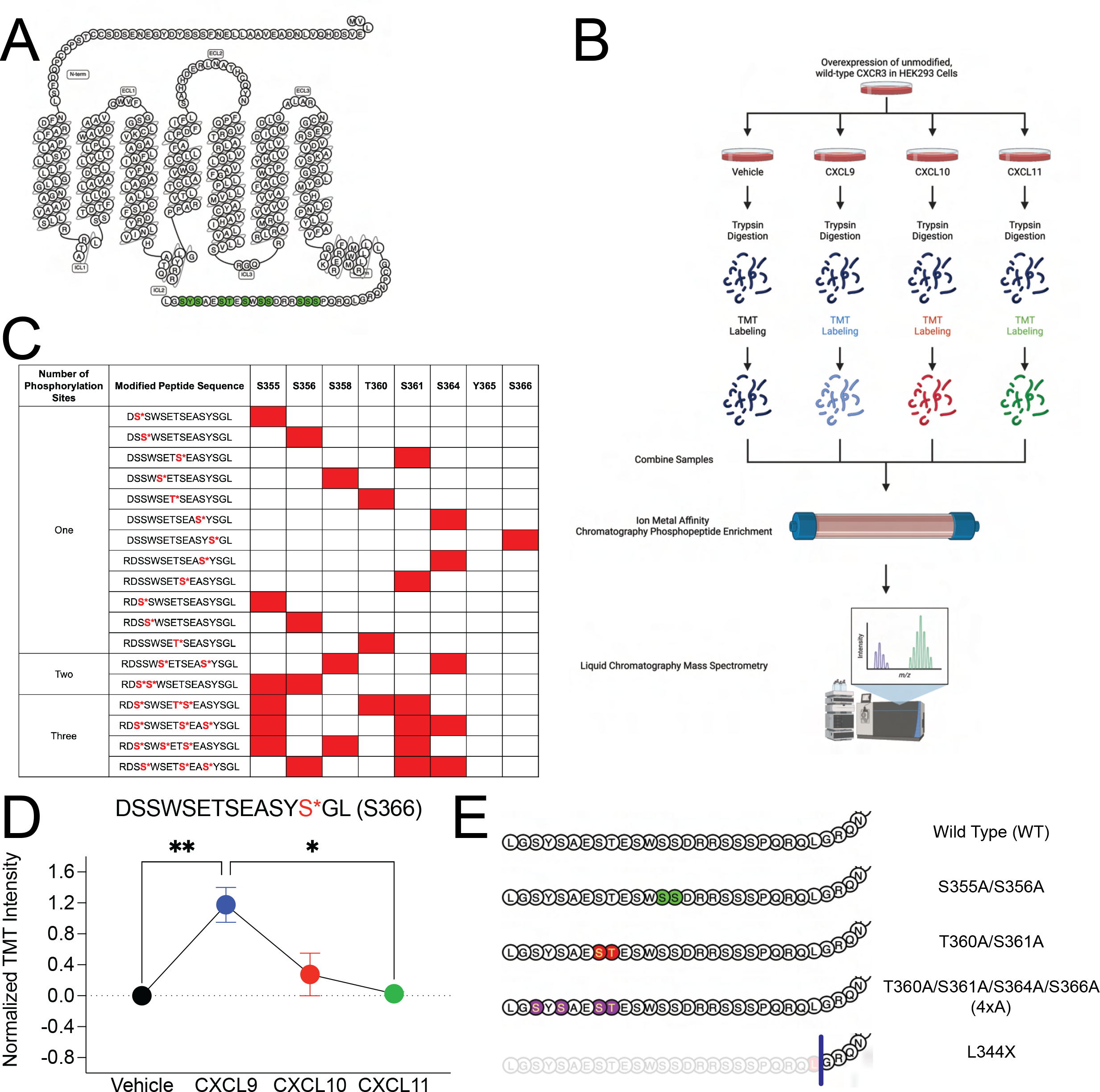
Detection of CXCR3 C-terminal phosphopeptides using mass spectrometry (**A**) Snake diagram of CXCR3 highlighting green putative C-terminal phosphorylation sites (S, T, and Y). (**B**) Schematic of experimental design of receptor phosphoproteomics experiment. (**C**) Singly, doubly, and triply phosphorylated CXCR3 C-terminal peptides identified through mass spectrometry. Identified phosphopeptides are noted in red. (**D**) Abundance of singly phosphorylated DSSWSETSEASYpSGL peptide measured in HEK293 cells following stimulation with vehicle control or CXCL9, CXCL10, or CXCL11 at 100 nM for 5 minutes. Mean ± SEM, n=2 technical replicates of 6 pooled biological replicates. (**E**) Diagram of designed CXCR3 phosphorylation-deficient receptor mutants of interest. *P<.05, by one-way ANOVA, Tukey’s post hoc analysis. See S1 for additional mass spectrometry data and signaling and expression data of CXCR3 phosphorylation deficient mutants.

We identified several specific phosphopeptides following chemokine treatment (**Figure 1C**). We detected that every putative serine or threonine phosphorylation site on the RDSSWSETSEASYSGL tryptic peptide could be phosphorylated, although the levels of these phosphopeptides differed depending on chemokine treatment. For example, the abundance of the singly phosphorylated peptide DSSWSETSEASYpSGL (S366) significantly increased following treatment with CXCL9 but did not change with CXCL10 or CXCL11, providing direct evidence that the chemokines encode distinct GPCR phosphorylation ensembles (**Figure 1D**). We additionally detected a decrease in abundance of singly phosphorylated peptides at S355, S356, and T360 following treatment with all chemokines (**Figure S1A-S1C**), consistent with a loss of some singly phosphorylated peptides following ligand treatment, as the ensemble of barcodes shift towards those that were multiply phosphorylated.

### Phosphorylation barcode ensembles direct G protein activation, β-arrestin recruitment, and receptor internalization

To study the effects of different phosphorylation barcode ensembles on cellular signaling, we screened a variety of phosphorylation-deficient CXCR3 mutants (**Figure S1D-S1G**), either serine/threonine to alanine mutants or truncation mutants, using G protein and β-arrestin assays. Based on this screen, we selected four phosphorylation deficient receptors to interrogate in detail (**Figure 1E**). These receptors maintained cell surface expression similar to wild-type CXCR3 (CXCR3-WT) (**Figure S1H**).

We first employed the TRUPATH bioluminescence resonance energy transfer (BRET) assay to assess G protein activation (Olsen et al., 2020) (**Figure 2A**). Most C-terminal mutations did not impact CXCR3 G protein activation, with similar profiles of CXCL11 and CXCL10 and reduced potency and Emax of CXCL9, as previously described (Smith et al., 2017) (**Figure 2B-2D**). We did observe a significant left shift in the EC50 of the truncation mutant CXCR3-L344X at CXCL10 and CXCL11, consistent with increased G protein signaling (**Figure 2C** and **2D**). When experiments were repeated in β-arrestin-1/2 CRISPR KO cells, CXCR3-WT potency was also left shifted and indistinguishable from the truncation mutant, consistent with CXCR3-L344X increased G protein signaling being due to a loss of β-arrestin-mediated desensitization (**Figure 2G**). Notably, we observed an approximately 50% decrease in G protein activation at the phosphorylation deficient mutant CXCR3- S355A/S356A (**Figure 2F**), which was not due to increased β-arrestin desensitization (**Figure S2A-S2D**), and was partially rescued using a phosphomimetic mutant, CXCR3-S355D/S356D (**Figure S2E-S2G**), consistent with receptor phosphorylation at specific sites impacting G protein activation.

**Figure 2:**
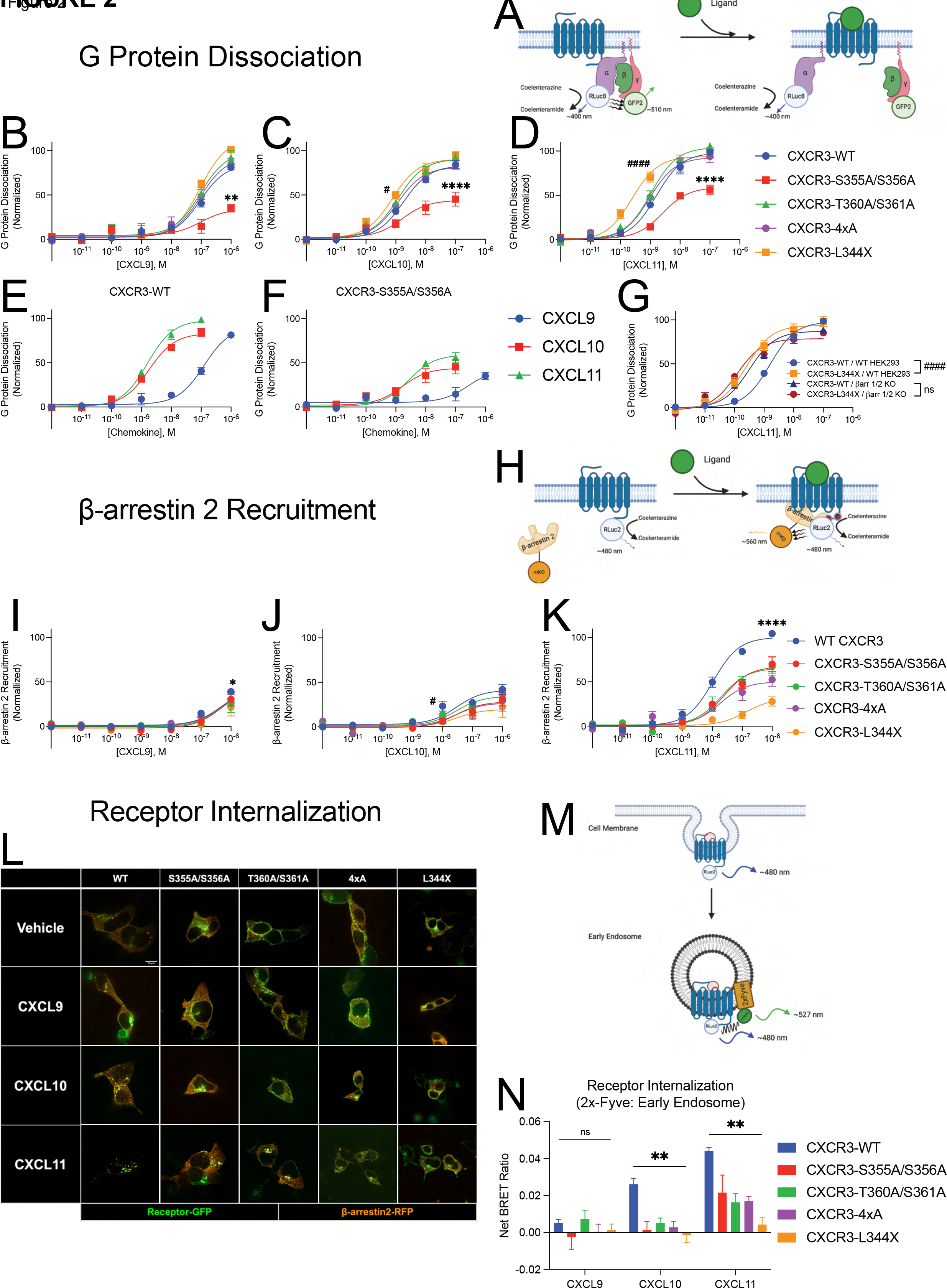
G protein dissociation, β-arrestin-2 recruitment, and receptor internalization of CXCR3 and receptor mutants (**A**) Schematic of TRUPATH assay to detect G protein dissociation following receptor stimulation using BRET (Olsen et al., 2020). (**B**, **C**, and **D**) G protein dissociation of receptors treated with listed chemokine in HEK293 cells. (**E** and **F**) G protein dissociation of CXCR3-WT and CXCR3-S355A/S356A in HEK293 cells. (**G**) G protein dissociation of CXCR3-WT and CXCR3-L344X in wild-type HEK293 cells (WT HEK293) and β-arrestin-1/2 knock out cells (βarr 1/2 KO). (**H**) Schematic of BRET assay to detect β-arrestin-2 recruitment to the receptor. (**I**, **J**, and **K**) β-arrestin-2 recruitment of receptors treated with listed chemokine in HEK293 cells. (**L**) Representative confocal microscopy images of HEK293 cells transfected with receptor-GFP and β-arrestin-2-RFP following treatment with vehicle control or the listed chemokine for 45 minutes. Images are representative of three biological replicates. (**M**) Schematic of BRET assay to detect receptor internalization to endosomes. (**N**) BRET data of receptor internalization following stimulation with the listed chemokine. Data are the average of BRET values from 20-30 minutes following ligand stimulation. For (**A-G**) TRUPATH assays, data shown are the mean ± SEM of BRET values 5 to 10 minutes following ligand stimulation, n=3. * denotes statistically significant differences between Emax of specified receptor and CXCR3-WT. # denotes statistically significant differences between EC50 of specified receptor and CXCR3-WT. For β-arrestin-2 recruitment, data shown are the mean ± SEM, n=3. *denotes statistically significant differences between EMax of CXCR3-WT and all other receptors at CXCL11, and of CXCR3-WT and CXCR3-4xA at CXCL9. # denotes statistically significant differences between EC50 of CXCR3-WT and CXCR3-S355A/S356A at CXCL10. For internalization BRET assays (**N**), data are the mean ± SEM, n=4. *P<.05 by two-way ANOVA, Dunnett’s post hoc testing between CXCR3-WT and all other receptor mutants. See S2 and S3 for further data assessing G protein dissociation, β-arrestin-2 recruitment, and receptor internalization.

We next examined β-arrestin recruitment (**Figure 2H**). Consistent with prior work, CXCL11 was significantly more effective in recruiting β-arrestin to CXCR3-WT compared to CXCL9 and CXCL10 (**Figure 2I- K**) (Colvin *et al*., 2004; Smith *et al*., 2017; Zheng *et al*., 2022). All phosphodeficient mutant receptors treated with CXCL11 demonstrated significantly less recruitment of β-arrestin when compared to CXCR3-WT (**Figure 2K**). In contrast, few mutant receptors treated with CXCL9 and CXCL10 demonstrated changes in β-arrestin recruitment relative to wild type (**Figure 2I-2J**). Differential β-arrestin recruitment was observed between chemokines at CXCR3-WT, CXCR3-S355A/S356A, and CXCR3-T360A/S361A (**Figure S2H-S2L**). In contrast, we observed no difference between chemokines in their ability to recruit β-arrestin to receptor mutants lacking the most putative C-terminal phosphorylation sites (CXCR3-4xA and CXCR3-L344X). This is consistent with CXCR3 C-terminal phosphorylation sites being critical for differences in β-arrestin recruitment between chemokines.

We next explored the impact of phosphodeficient CXCR3 receptors on β-arrestin function. β-arrestins are known to regulate GPCR endocytosis by interacting with the clathrin adaptor protein AP-2 (Ferguson et al., 1996; Kim and Benovic, 2002; Laporte et al., 1999). Therefore, we hypothesized that CXCR3 C-terminal mutations would impair receptor internalization. We used confocal microscopy to monitor CXCR3 and β-arrestin localization following chemokine treatment. CXCL11 promoted the translocation of CXCR3-WT and β-arrestin-2 to endosomes (**Figure 2L** and **Figure S3A**). CXCL10 also promoted CXCR3-WT:β-arrestin puncta, but not to the magnitude of CXCL11 (**Figure S3A**). CXCL9 did not promote either CXCR3-WT:β-arrestin puncta or receptor internalization **(Figure S3A)**. Consistent with our hypothesis, CXCL10 or CXCL11 treatment of phosphorylation-deficient CXCR3 mutants impaired internalization (**Figure 2L and S3B-S3E)**.

To further evaluate and quantify internalization, we utilized a BRET-based assay to measure receptor trafficking to early endosomes (**Figure 2M**). CXCL9 treatment did not induce significant CXCR3-WT endosomal trafficking (**Figure 2N**). While CXCL10 promoted receptor internalization, none of the phosphorylation-deficient mutants internalized after CXCL10 treatment. In contrast, CXCL11 treatment led to a different endosomal trafficking pattern at mutant receptors, with CXCR3-S355A/S356A, -T360A/S361A, and -4xA internalizing∼50% of the level of CXCR3-WT, while CXCR3-L344X did not internalize at all. We confirmed these findings with an orthogonal BRET assay to assess CXCR3 trafficking away from the plasma membrane following chemokine treatment (**Figure S3F-S3G**). These results are consistent with ligand- and receptor-specific effects on internalization: while removing selected phosphosites is sufficient to seemingly eliminate CXCL10-mediated internalization, removal is not sufficient to completely inhibit CXCL11-mediated internalization. In contrast, the receptor C-terminus is required for receptor internalization with CXCL10 and CXCL11, despite partial β-arrestin recruitment to the CXCR3-L344X mutant.

### GRK2 and GRK3 are differentially recruited to CXCR3 following ligand stimulation

We next investigated the kinases critical to differential CXCR3 phosphorylation barcode ensembles. While multiple kinases have been identified that phosphorylate GPCRs, the GRKs are established to be the primary drivers of GPCR phosphorylation (Gurevich and Gurevich, 2019; Komolov and Benovic, 2018; Tobin, 2008). There are seven identified GRK isoforms, of which GRK2, 3, 5, and 6 are ubiquitously expressed in mammalian tissues (Ribas et al., 2007). Because CXCR3 is primarily expressed on leukocytes, we investigated GRKs 2, 3, 5 and 6 recruitment to CXCR3 following chemokine treatment using a previously validated nanoBiT complementation assay (Inoue et al., 2019). We observed that GRK2 and GRK3 were recruited to all CXCR3 mutant receptors following chemokine treatment with similar kinetic profiles (**Figure 3A-3F**). In contrast, we did not observe appreciable recruitment of GRK5 or GRK6 to CXCR3-WT or mutant receptors, and confirmed it was not due to competition between GRK isoforms by demonstrating a lack of GRK5 and 6 recruitment in GRK2/3/5/6 knock out (KO) cells (Pandey et al., 2021a) (**Figure S4**).

**Figure 3:**
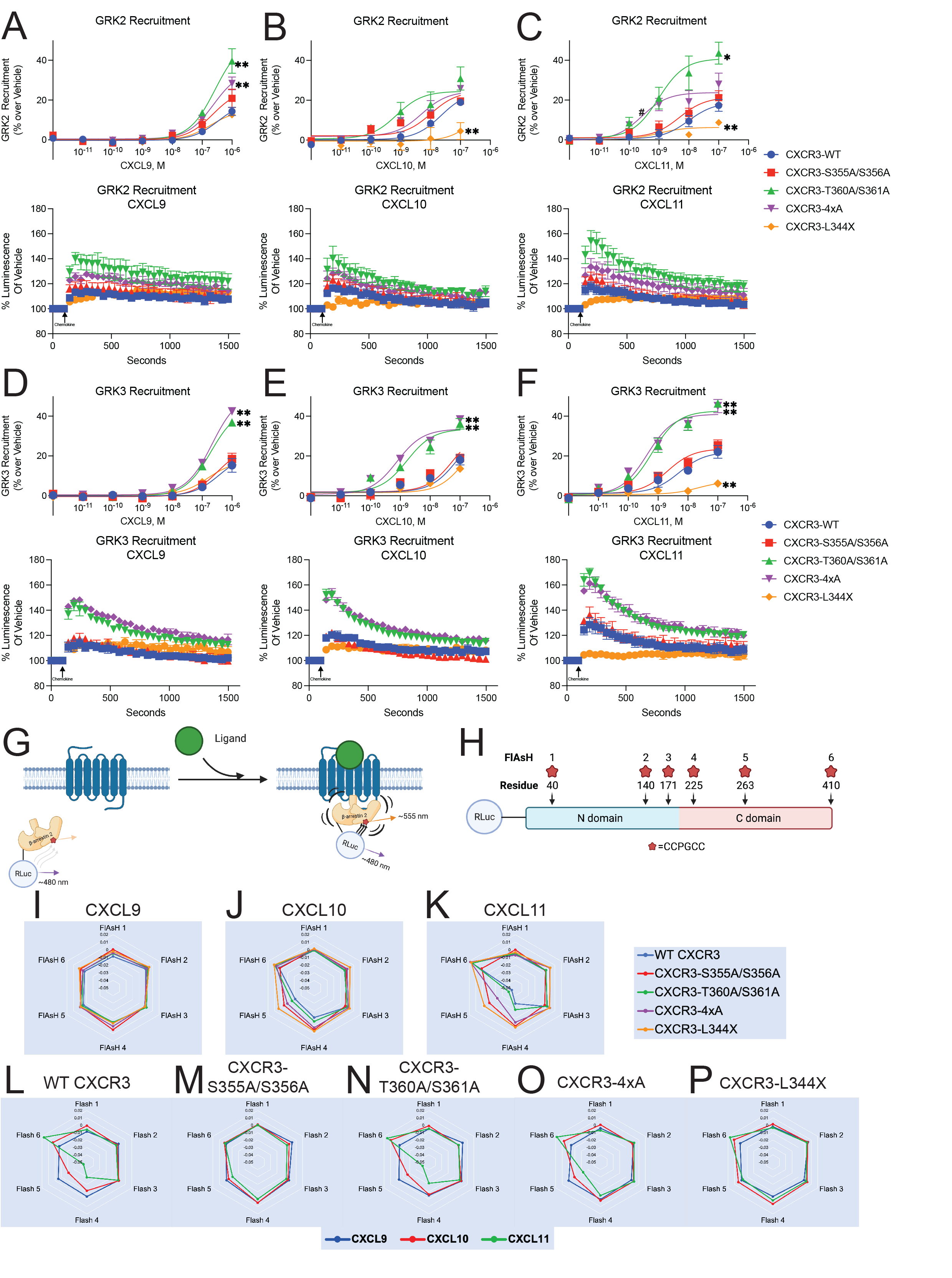
GRK Recruitment and β-arrestin-2 conformational dynamics Agonist dose-dependent data and kinetic data of maximum treatment dose of (**A-C**) GRK2 and (**D-F**) GRK3 recruitment to receptor as measured by a split nanoluciferase assay. Data are grouped by treatment condition. Mean ± SEM, n=3-4. (**G**) Schematic of FlAsH assay to detect β-arrestin-2 conformational dynamics following receptor stimulation using intramolecular BRET (Lee et al., 2016). (**H**) Location of N-terminal RLuc and CCPGCC FlAsH-EDT2 binding motifs on β-arrestin-2. (**I**-**K**) Radar plots of FlAsH 1-6 grouped by treatment. (**L-P**) Radar plots of FlAsH 1-6 grouped by receptor. Mean, n=5. For FlAsH BRET (**I-P**), data is the average of five consecutive reads taken approximately 10 minutes after the addition of ligand. See S4-S5 for additional GRK recruitment data and S6 for raw FlAsH data.

At CXCR3-WT, CXCL9, CXCL10, and CXCL11 demonstrated similar maximal recruitment of GRK2 and GRK3 (**Figure S5A** and **Figure S5L**). The effects of CXCR3 C-terminal mutations were variable (**Figure S5B-E** and **S5M-P**), with effects that were both chemokine- and receptor-dependent. At CXCR3-L344X, GRK2 and GRK3 recruitment was largely preserved with CXCL9 stimulation, but significantly reduced with CXCL10 or CXCL11 treatment (**Figure 3A-3C)**. Surprisingly, two phosphodeficient mutants enhanced GRK recruitment to the receptor. To investigate this, we generated the phosphomimetic mutants CXCR3-T360D/S361D and CXCR3- 4xD and found that they displayed decreased recruitment of GRK2 and GRK3, similar to that of CXCR3-WT (**Figure S5F-S5K** and **S5Q-S5V**). These results suggest that basal phosphorylation of specific residues in the C-terminus inhibit GRK recruitment. Together, these experiments demonstrate that GRK recruitment depends on both the specific C-terminal residue as well as the ligand used to activate the receptor.

### β-arrestin conformation is dependent on ligand identity and receptor phosphorylation status

We next investigated how chemokines modulate β-arrestin conformation. Previous work has shown that β-arrestins adopt multiple conformational states when engaged with the receptor core and C-terminus, and that these different states are important for β-arrestin-dependent signaling (Dwivedi-Agnihotri *et al*., 2020; Gurevich and Gurevich, 2004; He et al., 2021; Latorraca *et al*., 2020; Shukla et al., 2008; Xiao et al., 2004). We used a previously validated intramolecular fluorescent arsenical hairpin (FlAsH) BRET assay to assess β-arrestin conformation (**Figure 3G** and **3H**) (Lee et al., 2016) at all five mutant CXCR3 receptors treated with CXCL9, CXC10, or CXCL11. Data are presented as radar plots, enabling simultaneous visualization of all FlAsH biosensors at each receptor:ligand combination (**Figure 3I-3P**, conformation heat maps and bar charts corresponding to FlAsH signals are shown in **Figure S6A-S6F, S6G**). At CXCR3-WT, we found that CXCL9 did not induce a significant conformational change compared to vehicle, while both CXCL10 and CXCL11 promoted significant changes in the β-arrestin C-domain (FlAsH 4,5) and C-terminus (FlAsH 6) (**Figure S6G**). Minimal conformational changes were noted in the N-domain of β-arrestin (FlAsH 1,2).

Analyzing conformational signatures by chemokine, phosphorylation-deficient mutants had no significant effect on β-arrestin conformational signatures following treatment with CXCL9 (**Figure 3I**) but had significant effects on the conformations when stimulated with CXCL10 and CXCL11 (**Figure 3J** and **3K**). Analyzing conformational signatures by mutant, CXCR3-S355A/S356A abolished all chemokine-specific β-arrestin conformational signatures (**Figure 3M**). In contrast, CXCR3-T360A/S361A **(Figure 3N)** promoted a β-arrestin-2 conformational signature nearly identical to CXCR3-WT (**Figure 3L**). CXCR3-4xA had decreased conformational changes in the β-arrestin C-domain (FlAsH 4 and 5) compared to CXCR3-WT, but with preserved conformational changes at the C-terminus (FlAsH 6) (**Figure 3O**). At CXCR3-L344X, nearly all conformational differences between chemokines were lost, with only small differences observed in the β-arrestin C-terminus between chemokines (FlAsH 6) **(Figure 3P)**. These data suggest that, even in the absence of a C-terminus, the chemokines are still able to promote distinct β-arrestin-2 conformational signatures through the receptor core (**Figure S6G**). Phosphorylation sites in the C-terminus play a central role in determining β-arrestin 2 conformation, with sites such as S355 and S356 being critical for biased G protein activation, β-arrestin recruitment, and β-arrestin conformation.

These data further show that the conformational status of the N-domain (FlAsH 1,2) depends on the identity of the chemokine and receptor, but that these effects are largely independent of each other (**Figure S6A** and **S6B**). However, the totality of conformational data demonstrates that the conformational signature of the C- domain (FlAsH 4 and 5) and C-terminus (FlAsH 6) of β-arrestin is distinctively dependent on the combination, rather than additive effects of chemokine identity and CXCR3 phosphorylation status (**Figure S6D-S6F**).

### Molecular Dynamic Studies of β-arrestin

To better characterize the conformational changes of β-arrestin observed using FlAsH probes, we performed structural modeling and computer simulation. The exact location of probes 1-3 in the N-domain and probes 4-5 in the structured beta-sheets of the C-domain are highlighted in our structural model of β-arrestin-2 fused to RLuc **(Figure S7)**. As probe 6 is located within the distal C-tail, a highly flexible region which currently has not been crystallized, it is absent from our structural model and further analysis.

According to the BRET data, the signal from probes 1-3 (located in the N-domain) was similar in the presence of different chemokines (**Figure 3I-3K**) or C-tail mutations (**Figure 3L-3P**). This suggests that the relative position of the N-domain and RLuc to each other do not significantly change in those conditions. In contrast, we observed that probes 4 and 5 located in the C-domain are sensitive to different chemokines and C-tail mutations. This indicates that structural changes induced in β-arrestin 2 by the receptor and the type of chemokine ligand result in a significant positional change of the C-domain with respect to the RLuc-fused N-domain. Such observed conformational changes are likely the result of receptor-induced activation of β-arrestin 2. Interestingly, previous studies have highlighted that this activation of arrestin is linked to a twist of the C-terminal domain in respect to the N-domain (Chen et al., 2017; Dwivedi-Agnihotri et al., 2020; Latorraca et al., 2018; Shukla et al., 2013). This transition can be quantified using the interdomain rotation angle, with higher values of this angle being linked to more active-like conformations of β-arrestin and vice versa (**Figure S7**).

To investigate whether the interdomain twist correlates with the distance between probes 4 or 5 and the RLuc anchor point (Arg8), we monitored both descriptors in β-arrestin simulations starting from an active-like conformation (**Figure 4A**). As we did not include either the receptor or a C-tail in the system, such a setup allows β-arrestin to spontaneously inactivate (Latorraca et al., 2018) and to sample interdomain rotation angles from 20 (active-like state) to 0 degrees (inactive-like state). Importantly, our simulations confirm that there is indeed a correlation between the RLuc-probe 4/5 distances and the interdomain rotation angles (R=0.54 for probe 4 and R=0.65 for probe 5). This suggests that these probes are sensitive to the activation state of β-arrestin 2. In contrast, the distances for probes 1-3 in the N-domain did not show any correlation with the value of the interdomain twist.

**Figure 4:**
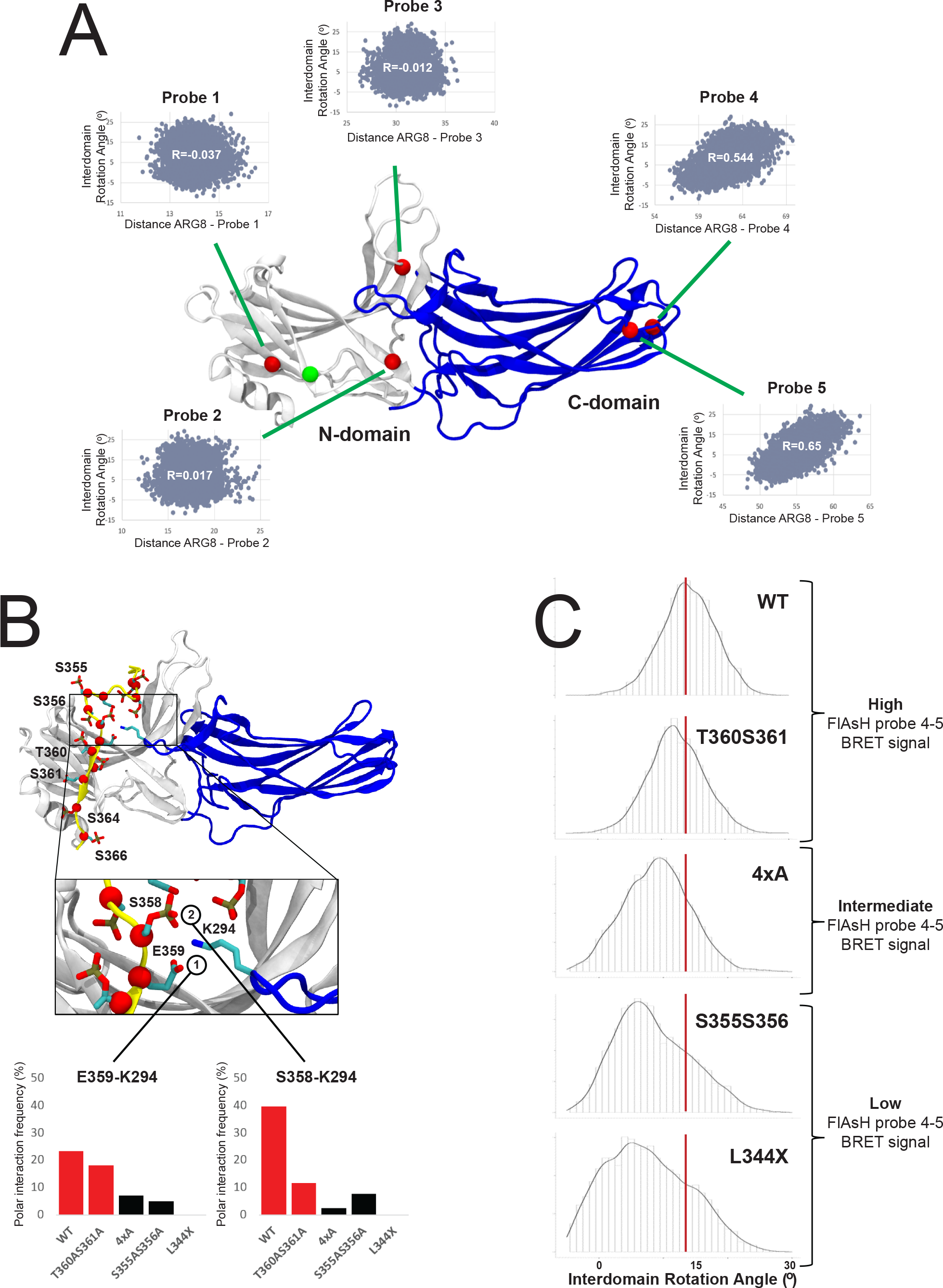
Impact of the phosphorylation pattern on β-arrestin-2 β-arrestin-2 conformational dynamics (**A**) Structural model of the construct used in the FlAsH BRET conformational assay. The positions of Probes 1- 5 are depicted as red spheres. Shown are the correlations between the distance of studied FlAsH probes to the RLuc domain and the interdomain rotation angle of β-arrestin 2. As the RLuc domain is absent in the simulated system, distance from the studied probes was approximated to a residue in the beginning of the N-terminal domain (the attachment point of the RLuc), depicted as a green sphere. **B**) The β-arrestin 2/WT-CXCR3 C-tail complex. Negatively charged residues (Asp, Glu or phosphorylated Ser and Thr) on the C-tail are depicted in licorice and their Cα atoms are highlighted with red spheres. Positions mutated within this study are labeled. The insert provides a detailed depiction of the lariat loop region of β-arrestin 2 (blue) and interactions with negatively charged residues of the C-tail. Bar charts demonstrate the stability of polar interactions between K294 of the lariat loop and S358 and E359 of the C-tail. Values for the WT and T360AS361A systems are colored in red. (**C**) Density plots depicting interdomain rotation angles assumed by β-arrestin-2 during MD simulations with C-tail mutants.

To further verify this finding, we simulated β-arrestin 2 in complex with each of the studied CXCR3 C-tail variants (**Figure 4B**) and monitored their interdomain rotation angles (**Figure 4C**). We found that the WT samples primarily conformations with a rotation angle of 14°. A similar ensemble of conformations was also observed for the T360A/S361A mutant (peak 11°). Interestingly, the 4xA mutant showed a reduction in rotation angle whereas the strongest shift towards low rotation angles was found for the S355A/S356A and L344X mutants (peaks at 6° and 4°). The order of adopted rotation angles is consistent with the magnitude of BRET signal for FlAsH probes 4 and 5, demonstrating that these probes are useful tools to approximate β-arrestin activation.

β-arrestin structural dynamics provides a potential explanation for the induced conformational differences by specific C-tail mutants. We found that in the WT receptor, two negatively charged residues present in the C-tail (phosphorylated S358 and E359) form a bifurcated interaction with the positively charged residue K295 located in the lariat loop of β-arrestin 2 (**Figure 4B**, blue region). We observed that in systems which explore more active-like conformations (e.g., WT and T360A/S361A), there were, on average, more interactions between the lariat loop and the C-tail in comparison to systems that explored more inactive-like states (e.g., 4xA, S355A/S356A, L344X) (**Figure 4C**). Importantly, these findings are consistent with previous studies that have demonstrated that polar interactions of the C-tail with the lariat loop are functionally important (Baidya et al., 2020) and promote active-like conformations of β-arrestin (Dwivedi-Agnihotri et al., 2020).

### Global LC-MS proteomic and phosphoproteomic analyses reveal substantial variation in intracellular signaling between CXCL9, CXCL10, and CXC11

To further understand the breadth of intracellular signaling promoted by CXCR3 chemokines, we interrogated the global proteome of HEK293 cells treated with CXCL9, CXCL10, or CXCL11 (**Figure S8A**). We successfully identified over 150,000 total peptides corresponding to approximately 11,000 proteins. Of these peptides, approximately 30,000 were also identified as phosphopeptides corresponding to approximately 5,500 unique phosphoproteins (**Figure S8B** and **S8C**). The majority of identified phosphosites were phosphoserines and phosphothreonines, with a high degree of reproducibility across replicates (**Figure S8D** and **S8E**). We identified approximately 1,500 phosphopeptides that underwent significant changes in abundance following chemokine treatment (**Figure 5A**). We then performed a clustering analysis of those phosphopeptides to uncover coregulated signaling pathways (Rigbolt et al., 2011) (**Figure S8F**). Certain signaling pathway clusters were similar between the three chemokines, while other clusters demonstrated significant differences between treatments (**Figure 5B**). Gene ontology term enrichment was performed on the significantly divergent phosphopeptides to assess the biological processes, molecular functions, and cellular compartments regulated by CXCR3 (**Figure 5C-5E**). These analyses reveal differential regulation of cellular transcription, post-translational modifications, cytoskeletal rearrangements, and cellular migration (among others) between chemokines. Additionally, the nucleus, cytoplasm, cytoskeleton, and cell-cell junctions were the most identified cellular compartments found in our gene ontology analysis. These data also show that CXCR3 chemokines do not signal in a purely redundant manner and display a striking degree of heterogeneity across signaling pathways associated with multiple cellular functions and compartments.

**Figure 5:**
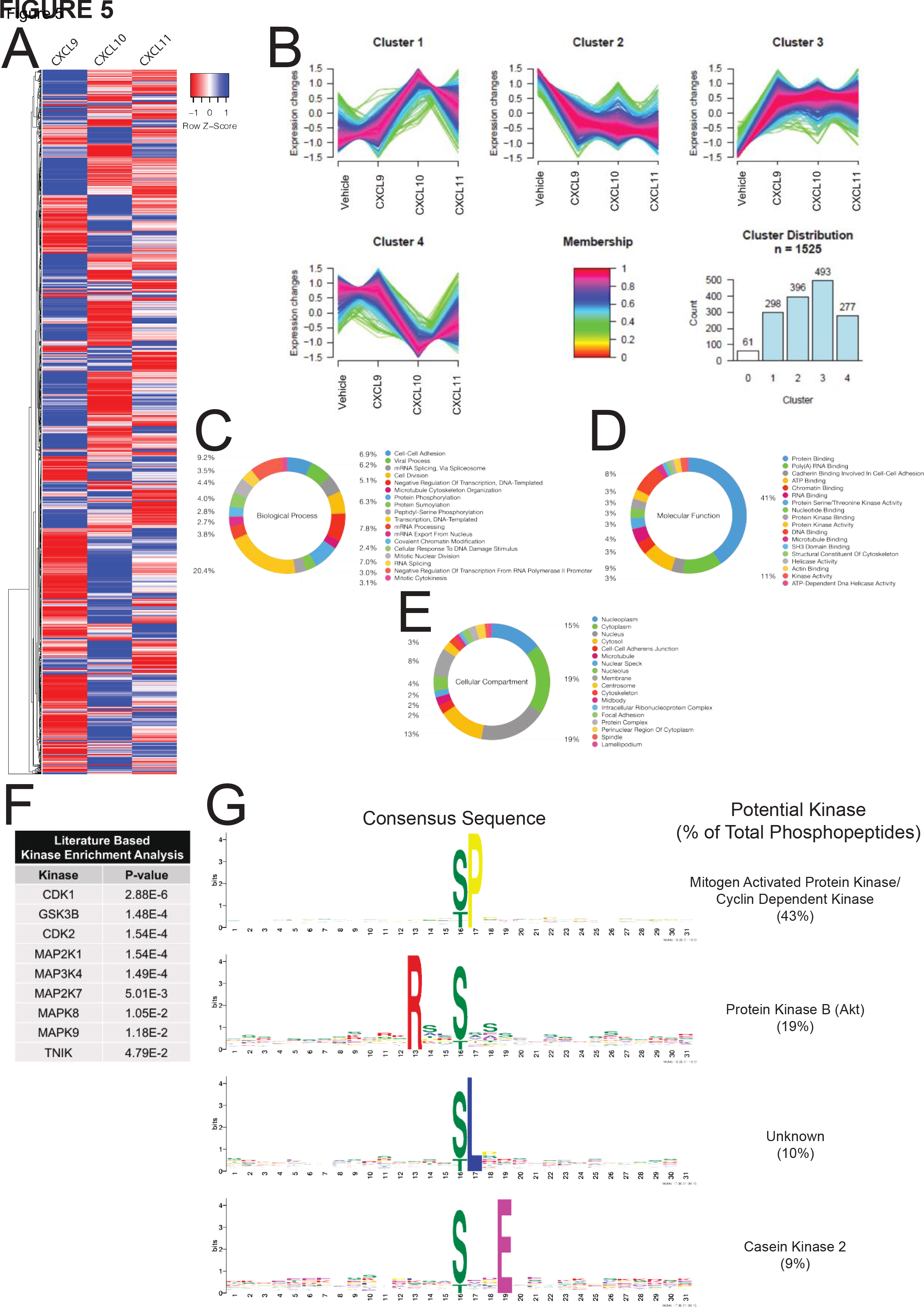
Characterization of the global phosphoproteome in HEK293 cells treated with endogenous CXCR3 agonist (**A**) HEK293 cells expressing CXCR3-WT were stimulated with vehicle control or 100 nM chemokine for five minutes. Heat map of statistically significant phosphopeptides normalized to vehicle control are shown. n=2 technical replicates of six pooled biological replicates. (**B**) Cluster analysis of significant phosphopeptides using GProX (Rigbolt *et al*., 2011). Cluster 0 is not shown for clarity due to low membership count. (**C-E**) Gene Ontology analysis of significant phosphopeptides as grouped by biological process, molecular function or cellular compartment, respectively. Percentiles demonstrate number of individual phosphopeptides present in each Gene Ontology Term. (**F**) Manually curated, literature-based kinase enrichment analysis to predict kinase activity based on significant phosphopeptides using Kinase Enrichment Analysis 2 (Lachmann and Ma’ayan, 2009). (**G**) Consensus sequences of significant phosphopeptides in the dataset as generated using MoMo from MeMe suite and identified kinases with listed consensus motif based on manual literature review (Keshava Prasad et al., 2009). See S7 for additional global phosphoproteomic data.

### Biased CXCR3 phosphorylation serves as a mechanism underlying differential regulation of the kinome

We next investigated the kinases responsible for generating chemokine-specific phosphorylation-dependent signaling networks. Kinase enrichment analyses (Lachmann and Ma’ayan, 2009) revealed that our dataset was largely enriched for phosphopeptides substrates targeted by cyclin dependent kinases (CDKs) and mitogen-activated protein kinases (MAPKs) (**Figure 5F**). We next used Modification Motifs, a motif-based sequence analysis tool (Bailey et al., 2006; Cheng et al., 2019), to identify enriched amino acid motifs flanking the phosphoserines and phosphothreonines that were differentially regulated in our dataset. Four major consensus sequences were identified: pS/pT-P which is a conserved target sequence of CDKs and MAPKs, R-X-X-pS/pT which is targeted by protein kinase B (Akt), pS/pT-L, and pS/pT-X-X-E which is targeted by casein kinase 2 (**Figure 5G**). These findings are consistent with previous studies demonstrating that CXCR3 activates the MAPK extracellular signal-related kinase (ERK) and Akt, among others (Bonacchi et al., 2001; Smith *et al*., 2018a), but also reveal unexplored CXCR3 signaling networks.

Next, we manually identified differentially phosphorylated kinases in our global phosphoproteomic data that are known to be regulated by GPCRs, or that were identified in bioinformatics analyses (**Figure 6A-6F**). The MAPKs ERK1, RAF1 and JNK, as well as SRC kinase family were phosphorylated in a chemokine-specific pattern, whereas BRAF and CSNK2 demonstrated similar phosphorylation patterns across CXCL9, CXCL10, and CXCL11. To understand if this biased regulation of the kinome is regulated by CXCR3 receptor phosphorylation, we studied ERK1/2 phosphorylation following chemokine treatment of cells expressing either CXCR3-WT or a phosphodeficient CXCR3 mutant (**Figure 6G-6J**). At CXCR3-WT, we saw a significant increase in phosphorylated ERK1/2 (pERK), consistent with our mass spectrometry results (**Figure S9A**). At 5 minutes, we observed a maximum increase in pERK levels over CXCR3-WT at CXCR3-4xA and CXCR3-L344X when stimulated with CXCL10 and CXCL11, but not CXCL9. At 30 and 60 minutes, pERK levels declined, consistent with prior observations at other GPCRs (Luttrell et al., 2018). This differential phosphorylation of ERK1/2 by the three chemokines was observed at all mutant CXCR3 receptors, including CXCR3-L344X (**Figure S9**).

**Figure 6:**
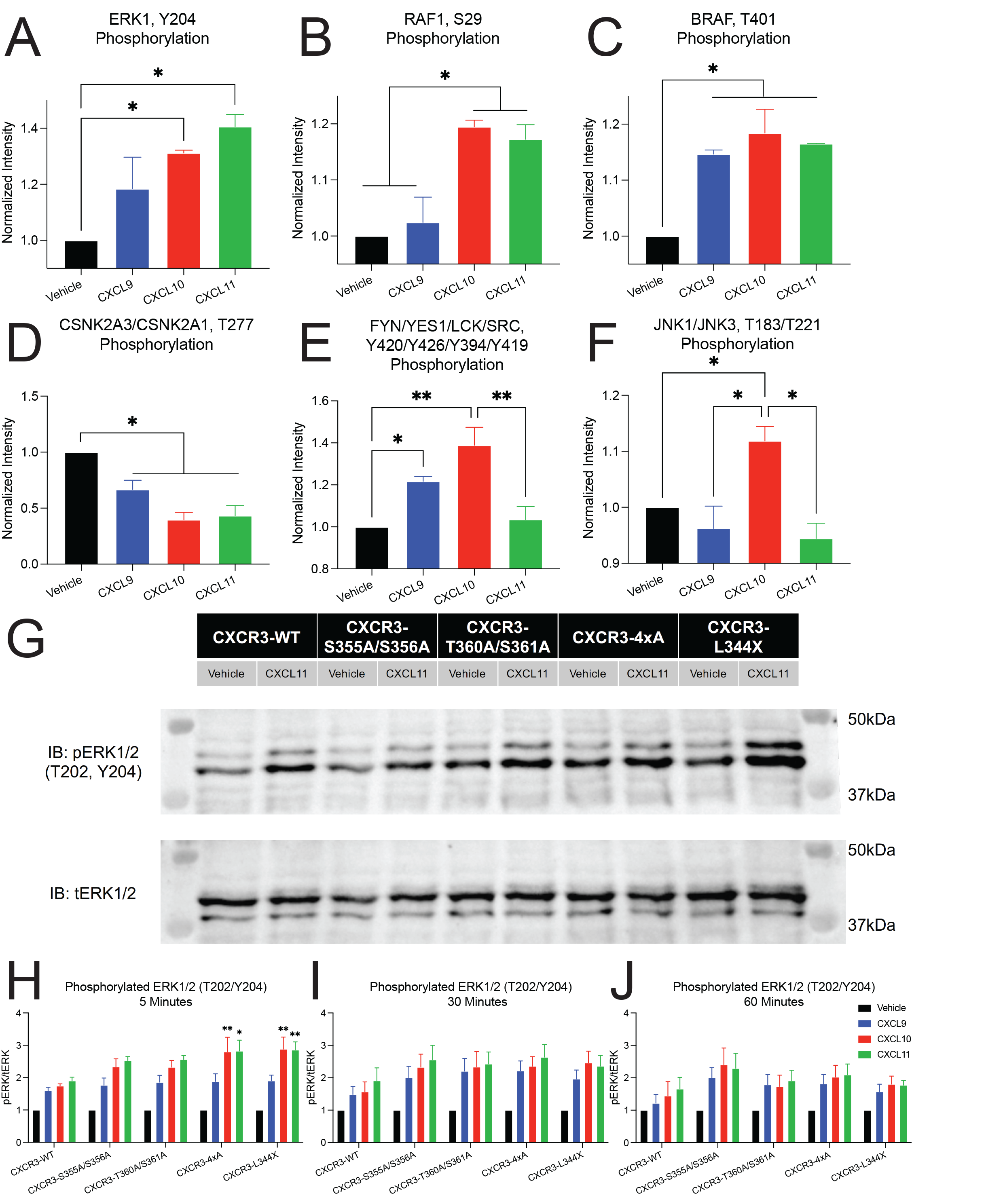
Differential regulation of kinases by biased ligands and phosphodeficient receptors Biased phosphorylation of various kinases identified from the global phosphoproteomics data including (**A**) ERK1, (**B**) RAF1, (**C**) BRAF, (**D**) Casein kinase 2 (CSNK2A3/CSNK2A1), (**E**) Src family of protein tyrosine kinases (FYN/YES1/LCK/SRC), and (**F**) JNKs (JNK1/JNK3). Data is normalized to vehicle treatment and n=2 technical replicates of six pooled biological replicates. Mean ± SEM. *P<.05 by one-way ANOVA, Tukey’s post hoc testing. (**G**) Representative western blot of phosphorylated ERK1/2 (pERK 1/2) and total ERK1/2 (tERK 1/2) following stimulation with vehicle control or 100 nM of CXCL11 for five minutes. (**H-J**) Quantification of western blots of pERK1/2 levels at 5, 30, and 60 minutes. Mean ± SEM, n=4. *P<.05 by two-way ANOVA, Dunnet’s post hoc testing denotes comparisons between a specific ligand/receptor combination to the same ligand at CXCR3- WT. See S8 for quantification of western blots grouped by receptor.

### T cell chemotaxis is regulated by biased CXCR3 phosphorylation barcode ensembles

We last investigated if the biased chemokine signaling pathways observed in HEK293 cells impact physiologically relevant cellular functions. Given that CXCR3 plays a central role in T cell function, we interrogated the effect of CXCR3 phosphorylation barcodes on T cell chemotaxis. We first used CRISPR/Cas9 to knock out the endogenous CXCR3 in Jurkat cells (an immortalized human T lymphocyte cell line), generating CXCR3 knockout (CXCR3-KO) Jurkats. We rescued CXCR3 receptors of interest (WT and mutants) with lentiviral constructs to generate stably expressing CXCR3+ Jurkat cell lines (**Figure 7A**). We confirmed similar receptor expression levels between WT and mutant CXCR3+ Jurkat lines (**Figure 7B**).

**Figure 7:**
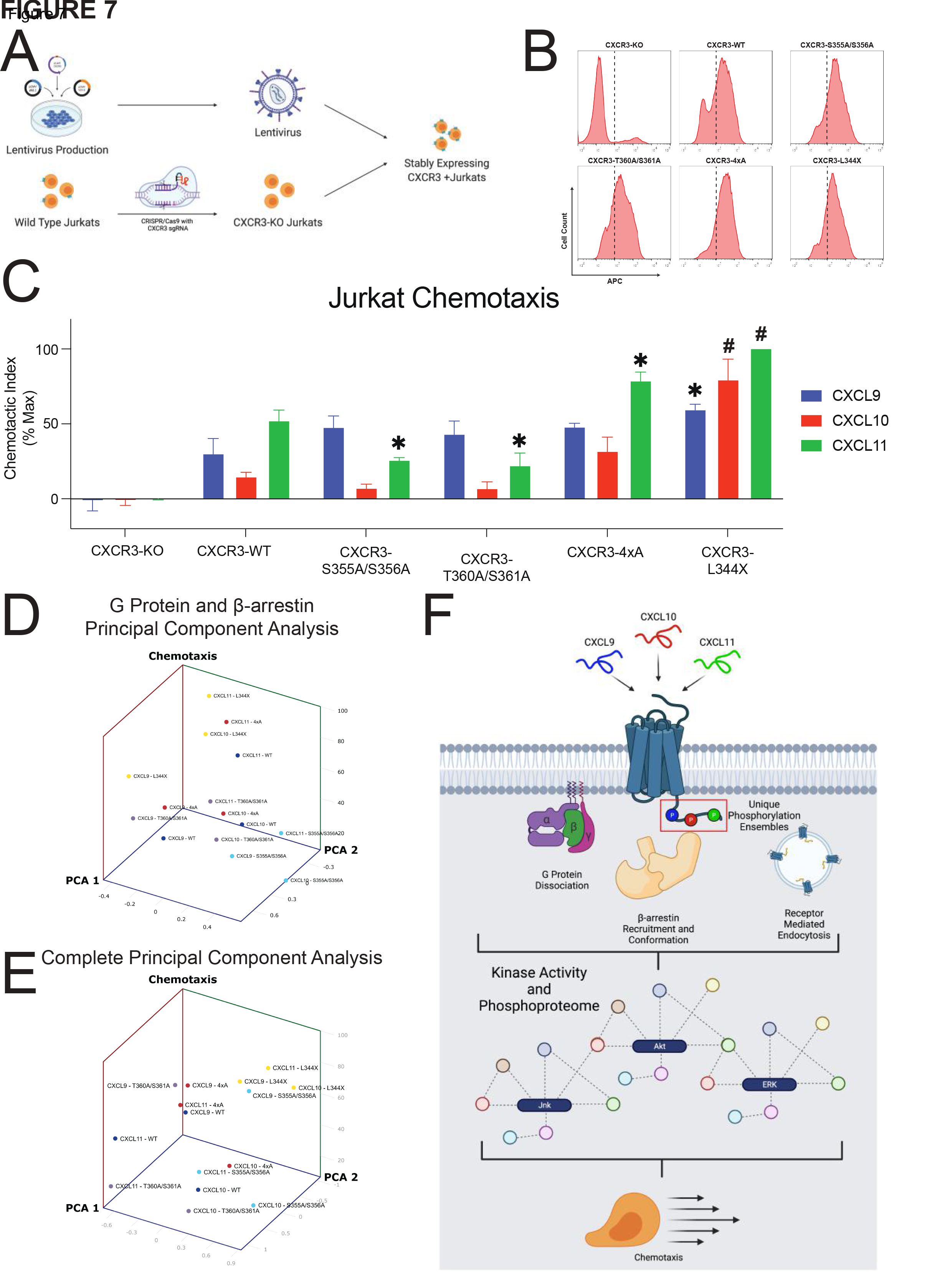
Jurkat chemotaxis and model of the phosphorylation barcode (**A**) Schematic of lentiviral production carrying cDNA for CXCR3-WT or one of the four receptor mutants, generation of CXCR3-KO Jurkats using CRISPR/Cas9, and creation of stably expressing CXCR3 Jurkats. (**B**) Surface expression of CXCR3-KO Jurkats or five various Jurkat cell lines transduced with lentivirus carrying the listed receptor cDNA as measured with flow cytometry. Dotted line denotes a fluorescence intensity of 10^2^. For transduced cells, cells with a fluorescence intensity greater than 10^2^ were sorted for chemotaxis experiments. (**C**) Jurkat chemotaxis for each receptor/ligand combination. Mean ± SEM, n=4. *P<.05 by two-way ANOVA, Tukey’s post hoc testing denotes comparisons between a specific ligand/receptor combination to the same ligand at CXCR3-WT. (**D**) Principal Component Analysis of G Protein activation and β-arrestin-2 recruitment versus chemotaxis. (**E**) Principal Component Analysis of G Protein activation, β-arrestin-2 recruitment, GRK2 and GRK3 recruitment, and FlAsH versus chemotaxis. See S9 for chemotaxis data grouped by receptor and univariate analyses. (**F**) Working model for biased ligand generation of unique barcode ensembles which differentially regulate G protein signaling, β-arrestin recruitment and conformation, receptor endocytosis, kinase activity, the global phosphoproteome, and cellular functions such as chemotaxis.

We then performed chemotaxis assays with these cell lines. Due to the promiscuous nature of the chemokine system, we first confirmed that CXCR3-KO Jurkats exhibit no measurable chemotaxis compared to vehicle treatment (**Figure 7C**), demonstrating that the observed chemotactic response is mediated by CXCR3 and not by other chemokine receptors. CXCR3+ Jurkat cells migrated with different chemotactic indices to CXCL9, CXCL10, or CXCL11, consistent with a biased response across chemokines. Statistically, there were effects induced both by ligand and by receptor (**Figure S10A-S10F**). We observed a slight but significant decrease in chemotactic function at CXCR3-S355A/S356A and CXCR3-T360A/S361A with CXCL11, but not with CXCL9 nor CXCL10. Conversely, we observed a significant increase in chemotaxis with CXCL11 at CXCR3- 4xA and CXCR3-L344X, although with different patterns. While chemotaxis at CXCR3-4xA displayed the same pattern by chemokine as CXCR3-WT (CXCL11 > CXCL9 > CXCL10), chemotaxis at CXCR3-L344X displayed only minor differences between all chemokines, although all displayed significantly more chemotaxis than at CXCR3-WT.

### Associating T cell chemotaxis with transducer efficacy

The biased pattern of chemotaxis at all receptors except L344X was different than that observed at proximal GPCR signaling assays, i.e., G protein activation and β-arrestin recruitment. To ascertain if G protein or β-arrestin signaling was directly related to chemotactic function, we performed univariate linear regressions on these data and found no significant linear relationship between G protein or β-arrestin signaling efficacy and chemotactic function (**Figure S10G-S10H**). We then performed a principal component analysis of G protein signaling and β-arrestin signaling versus chemotactic function and similarly found no obvious clustering of data by ligand or receptor (**Figure 7D**). A univariate linear regression of MAPK activation versus chemotactic function did demonstrate a significant positive linear relationship (**Figure S10I**). We then performed a second principal component analysis of all major assays conducted in this study and were able to demonstrate clustering of the chemokines at CXCR3-L344X (**Figure 7E**). These analyses demonstrated that G protein and β-arrestin signaling alone or together do not comprehensively describe the observed variance in our functional assays. Further addition of other signaling data (GRK recruitment, FlAsH conformational assays) moderately enhanced our ability to describe the variance in cellular chemotaxis, however, only at CXCR3-L344X. For receptors with a C-terminal tail, their chemotactic profiles did not cluster after PCA analysis, consistent with the C-terminus contributing to a biased response even when differences in transducer coupling are accounted for. Together, these data support a working model in which biased chemokines promote bias through the receptor core and through different CXCR3 phosphorylation barcode ensembles that regulate both proximal and distal aspects of GPCR signaling that impact T cell chemotaxis in a complex fashion (**Figure 7F**).

## DISCUSSION

Here we report how different chemokines for the same receptor direct distinct signaling pathways. We conclusively demonstrate that the endogenous chemokines of CXCR3 have biased patterns of signaling and are nonredundant in their activation of different intracellular kinase cascades and chemotactic profiles. Signal initiation through G proteins and β-arrestins, well-conserved effectors across the GPCR superfamily, is directed by CXCR3 C-terminus phosphorylation, whereby the different chemokines encode distinct phosphorylation ensembles. Disrupting discrete CXCR3 phosphorylation patterns interfered with signaling downstream of certain CXCR3 chemokines, but not others, depending on the phosphorylation site. Disrupting certain phosphosites also altered T cell function as assessed by chemotaxis, and this complex physiological output could not be entirely defined by the activity of proximal G protein or β-arrestin transducers alone.

Using multiple high-resolution mass spectrometry approaches, we found that different chemokines promoted different CXCR3 phosphorylation barcode ensembles. Limitations of mass spectrometry-based approaches in studying the phosphorylation of transmembrane receptors include their relative low abundance, difficulty in isolation, and sample handling demands. To overcome these challenges, we incorporated and validated a combinatorial phosphopeptide library with heavy isotope-labeled reference standards (Tsai et al., 2019), allowing us to simultaneously analyze wild-type, untagged CXCR3 under different chemokine treatment conditions. We found that perturbations in specific phosphorylation patterns impact proximal and distal aspects of GPCR signaling, as well as chemotaxis. At GPCRs more broadly, there is limited work investigating the phosphorylation patterns generated by endogenous ligands (Busillo et al., 2010), as most studies have relied on synthetic ligands (Butcher et al., 2011; Miess et al., 2018; Nobles et al., 2011). In addition, there is a desire to develop biased therapeutics which preferentially activate signaling pathways to increase therapeutic efficacy while simultaneously decreasing side effects, and our findings could provide an initial methodology to screen ligands for a desired physiological output. Our results demonstrate that the GPCR C-terminus is critically important in the regulation of proximal signaling effectors, and that the final cellular phenotype is dependent on the integration of multiple signaling pathways downstream of these interactions.

We found that both the receptor core and distinct phosphorylation patterns in the tail contribute to the allosteric regulation of β-arrestin conformation. β-arrestins can engage GPCRs through independent interactions with the receptor core and C-terminus (Cahill et al., 2017; Eichel et al., 2018; Kahsai et al., 2018; Latorraca *et al*., 2018). We found that all chemokine agonists similarly recruited β-arrestin to the receptor core in the absence of a C-terminus but maintained the ability to promote different β-arrestin conformations. Additionally, although β- arrestin-2 could still recruit to CXCR3-L344X, the receptor did not internalize, highlighting the importance of the β-arrestin-2 interaction with the receptor C-terminus in promoting receptor internalization as previously described (Cahill et al., 2017). Our findings agree with recent studies demonstrating that not all phosphorylation sites on a GPCR C-terminus impact β-arrestin recruitment and function (Dwivedi-Agnihotri et al., 2020; Latorraca et al., 2020). PCA analysis of signaling and chemotaxis data support a model in which chemokines promote bias through both the receptor core and CXCR3 phosphorylation barcode ensembles that regulate both proximal and distal aspects of GPCR signaling.

While previous studies demonstrated that certain C-terminal phosphorylation sites are involved in β- arrestin conformation, many of these studies have been limited to *in vitro* and *in silico* methods (Sente et al., 2018; Zhou et al., 2017). Here, we demonstrate in a cellular context that the β-arrestin conformation formed at a GPCR is dependent on the specific combination of both the ligand and the receptor phosphorylation pattern. Importantly, the conformational diversity seen in the C-domain and C-terminus of β-arrestin cannot be explained simply through the additive effects of ligand and receptor identity. Rather, the unique interaction between the ligand and receptor phosphorylation pattern ultimately promotes β-arrestin to adopt a specific ensemble of conformations, highlighting the complex structural diversity a single GPCR can impose upon proximal effector proteins like β-arrestin. Modeling and molecular dynamics simulations suggest that β-arrestin conformations vary in the degree of interdomain rotation between the N- and C-domains. This motion has been previously described to be a crucial step in β-arrestin activation (Latorraca *et al*., 2018). Our results show that certain chemokines and C-tail mutants shift the conformational equilibrium of β-arrestin towards active-like conformations. Furthermore, a more detailed analysis suggests that a specific pattern of interaction of the receptor C-tail with the lariat loop region of β-arrestin contributes towards this transition.

This work also provides a comprehensive assessment of the roles biased agonists and receptor phosphorylation serve in directing downstream signaling. Not only did we observed that chemokines induce distinct phosphoproteomic signaling profiles through a single receptor, but we also demonstrate how specific changes in CXCR3 phosphorylation barcodes impact the biased regulation of the phosphoproteome and MAPK signaling. We identified a relationship between MAPK activation and chemotactic function, even though these assays were performed in different cell types, consistent with previous studies (Shahabuddin et al., 2006; Sun et al., 2002). Our data suggests that a systems-level approach integrating upstream and downstream signaling effectors will be critical to the development of novel therapeutics with a desired phenotype, rather than an approach that relies solely on specific proximal transducers. Because protein-protein interfaces are frequent pharmacologic targets and commonly regulated via phosphorylation, this investigative framework extends to many other domains of pharmacology and cellular signaling (Stevers et al., 2018; Watanabe and Osada, 2012). Therefore, this study has important implications in understanding the pluridimensional efficacy of the chemokine system, the GPCR superfamily, and all receptors more broadly.

Our findings prompt many avenues for future study. Importantly, there are technical limitations that must be overcome to better determine the abundance of highly phosphorylated C-terminal peptides. Accurate determination of the stoichiometry of physiologically relevant phosphorylation barcodes is critical to understanding how these ensembles direct GPCR effector function under native conditions. Additionally, further work is needed to elucidate the detailed mechanism underlying the generation of these barcode ensembles – while we provide evidence demonstrating biased interactions of GRKs with CXCR3, it remains unclear how these ligands target the GRKs and other kinases to specific locations within the C-terminus and receptor core. Notably, there is heterogenous expression of the GRKs and other kinases throughout the human body; therefore, it is pertinent to understand how receptor phosphorylation may change depending on the effectors present to interact with a GPCR (Sato et al., 2015). Also, while there is evidence of specific signaling cascades directly dependent on G protein or β-arrestin activation, more complex cellular phenotypes are likely dependent on the combination of these and other GPCR signaling partners. For example, there is burgeoning evidence of GPCR signaling pathways that extend beyond that canonical G protein versus β-arrestin paradigm, specifically, those that integrate these pathways together (Smith et al., 2021). Using systems-level approaches to characterize these processes will be critical to understanding the coordination of signaling through different GPCR transducers.

While it was a long-held belief that signaling in the chemokine system was redundant (Mantovani, 1999), we conclusively demonstrate that signaling through the three endogenous chemokine agonists of CXCR3, CXCL9, CXC10, and CXCL11 is not redundant. These three chemokines (1) encode distinct receptor phosphorylation patterns, (2) promote strikingly divergent signaling profiles as assessed by ∼30,000 phosphopeptides corresponding to ∼5,500 unique phosphoproteins, and (3) promote distinct phosphosite-dependent physiological effects as assessed by chemotaxis. We have previously shown in a mouse model of contact hypersensitivity that a β-arrestin-biased CXCR3 agonist can increase inflammation whereas a G protein-biased CXCR3 agonist did not (Smith *et al*., 2018a), further supporting the physiological relevance of biased signaling at CXCR3. Additionally, T cells derived from β-arrestin-2 KO mice demonstrate impaired chemotactic response in the presence of either a β-arrestin-biased or G protein biased CXCR3 agonist (Smith *et al*., 2018a). Taken as a whole, our findings suggest that cellular functions such chemotaxis are not merely encoded by the amount of β-arrestin recruited to the receptor, rather, it is influenced by specific β-arrestin conformations induced by a receptor (Ge et al., 2003; Lin et al., 2018; Yang *et al*., 2015). The non-redundant nature of chemokine signaling at CXCR3 likely applies to the remainder of the chemokine system, although further work is necessary to confirm this hypothesis.

## ACKNOWLEDGEMENTS

We thank R.J. Lefkowitz (Duke University, USA) for guidance, mentorship, and thoughtful feedback throughout this work; N. Nazo for laboratory assistance; R. Shammas for assistance with FlAsH experiments. Funding: This work was supported by T32GM007171 (D.S.E.), the Duke Medical Scientist Training Program (D.S.E.), AHA 20PRE35120592 (D.S.E.), 1R01GM122798 (S.R.), K08HL114643 (S.R.), Burroughs Wellcome Career Award for Medical Scientists (S.R.), National Center of Science, Poland - 2017/27/N/NZ2/02571 (T.M.S), P41GM103493 (R.D.S.), 1R01GM139858 (T.S.). A.I. was funded by Japan Society for the Promotion of Science (JSPS) KAKENHI grants 21H04791, 21H05113, JPJSBP120213501 and JPJSBP120218801; FOREST Program JPMJFR215T and JST Moonshot Research and Development Program JPMJMS2023 from Japan Science and Technology Agency (JST); The Uehara Memorial Foundation; and Daiichi Sankyo Foundation of Life Science. Work was performed in the Environmental Molecular Sciences Laboratory, a U. S. Department of Energy Office of Biological and Environmental Research national scientific user facility located at Pacific Northwest National Laboratory in Richland, Washington. Pacific Northwest National Laboratory is operated by Battelle for the U.S. Department of Energy under Contract No. DE-AC05-76RLO 1830.

## AUTHOR CONTRIBUTIONS

Conceptualization, D.S.E., J.S.S., and S.R; Methodology, D.S.E., J.S.S., J.M.J, R.D.S., S.R., T.M.S., and J.D.S.; Investigation, D.S.E., J.S.S., C.H., N.B., J.G., T.S., C.T., N.K., C.D.N., A.M.M., T.M.S, K.K., I.C., K.Z., A.W., P.A., N.M.K., O.H.; Resources, K.K.,A.I.; Writing - Original Draft, D.S.E., J.S.S., and S.R.; Writing – Reviewing & Editing: D.S.E., J.S.S., and S.R, Visualization: D.S.E., J.S.S., and S.R; Supervision and Funding Acquisition, S.R.

## INCLUSION AND DIVERSITY

One or more of the authors of this paper self-identifies as an underrepresented ethnic minority in science. While citing references scientifically relevant for this work, we also actively worked to promote gender balance in our references list.

## STAR Methods

### RESOURCE AVAILABILITY

#### Lead Contact

Further information and requests for resources and reagents should be directed to and will be fulfilled by the lead contact, Sudarshan Rajagopal.

#### Materials Availability

All plasmids generated in this study will be distributed upon request.

#### Data and Code Availability

The RAW MS data and the identified results from Maxquant have been deposited in Japan ProteOme STandard Repository (jPOST: https://repository.jpostdb.org/) (Watanabe et al., 2021). The accession codes: JPST001599 for jPOST and PXD034033 for ProteomeXchange. The access link is https://repository.jpostdb.org/preview/1101419412628c1a4318aa7 and access key is 6844. Molecular dynamics simulations have been deposited in GPCRmd (https://submission.gpcrmd.org/dynadb/publications/1485/) with the ID 1485.

### EXPERIMENTAL MODEL AND SUBJECT DETAILS

#### Bacterial strains

XL-10 Gold ultracompetent E. coli (Agilent) were used to express all constructs used in this manuscript.

#### Cell Lines

Human Embryonic Kidney (HEK293, β-arrestin 1/2 knockout) cells were grown in minimum essential media (MEM) supplemented with 10% fetal bovine serum (FBS) and 1% penicillin/streptomycin at 37°C and 5% CO2. β-arrestin ½ KO HEK293 cells and GRK 2/3/5/6 KO HEK293 cells were provided by Asuka Inoue and validated as previously described (Alvarez-Curto et al., 2016; Pandey *et al*., 2021a). Jurkat cells were cultured in RPMI 1640 supplemented with 10% FBS and 1% penicillin/streptomycin at 37°C and 5% CO2.

### METHOD DETAILS

#### Cell culture and transfection

Human embryonic kidney cells (HEK293, GRK 2/3/5/6 knockout, β-arrestin 1/2 knockout) were maintained at 37°C and 5% CO2, in minimum essential medium supplemented with 1% penicillin/streptomycin and 10% fetal bovine serum (FBS). For BRET and luminescence studies, HEK293 cells were transiently transfected via an optimized calcium-phosphate protocol as previously described (Pack et al., 2018). In the calcium phosphate transfection method, cell culture media was replaced 30 minutes prior to transfection. Plasmids were suspended in water, and calcium chloride was added to the plasmid constructs to a final concentration of 250 µM. An equal volume of 2x HEPES-buffered saline solution (10 mM D-Glucose, 40 mM HEPES, 10 mM potassium chloride, 270 mM sodium chloride, 1.5 mM disodium hydrogen phosphate dihydrate) was added to the solution, allowed to incubate for two minutes, and subsequently added to the cells. For mass spectrometry studies and confocal microcopy, constructs were overexpressed in HEK293 cells using FuGENE 6 according to the manufacturer’s instructions (Promega, Madison, WI). For TGF-α shedding assay cells, were transiently transfected using Lipofectamine 2000 according to the manufacturer’s instructions (Thermo Fisher Scientific).

#### Generation of constructs

Cloning of constructs was performed using conventional techniques such as restriction enzyme and ligation methods. CXCR3 C-terminal phosphomutant constructs were generated using a QuikChange Lightening Mutagenesis Kit (Agilent, Santa Clara, CA). Linkers between the fluorescent proteins or luciferases and the cDNAs for receptors, transducers, kinases, or other proteins were flexible and ranged between 2 and 17 amino acids.

#### Cell lysis and protein extractions

For protein extraction, cell pellets were resuspended in cell lysis buffer (100 mM NH4HCO3, pH 8.0, 8 M urea, 75 mM sodium chloride (NaCl), 10 mM sodium fluoride (NaF), 1% phosphatase inhibitor cocktail 2 (Sigma P 5726), 1% phosphatase inhibitor cocktail 3 (Sigma P 0044), pH 8.0) and sonicated in an ice-bath for 3 mins followed by homogenization using a hand-held SpiralPestle™ and MicroTube Homogenizer (BioSpec products, Bartlesville, OK) on ice until complete visual homogenization was achieved. Cell lysates were centrifuged, and the protein concentrations were measured with a Pierce BCA protein assay (Thermo Fisher Scientific). Proteins were reduced with 5 mM dithiothreitol for one hour at 37°C and subsequently alkylated with 20 mM iodoacetamide for one hour at 25°C in the dark. Samples were diluted 1:8 with 50 mM NH4HCO3 and digested with sequencing-grade modified trypsin (Promega, V5117) at a 1:50 enzyme-to-substrate ratio. After three hours of digestion at 37°C, the digested samples were acidified with 100% formic acid (FA) to 1% of the final concentration of FA and centrifuged for 15 minutes at 1,500 *×g* at 4°C before transferring samples into new tubes leaving the resulting pellet behind. Digested samples were desalted using a 4-probe positive pressure Gilson GX-274 ASPEC™ system (Gilson Inc., Middleton, WI) with Discovery C18 100 mg/1 mL solid phase extraction tubes (Supelco, St.Louis, MO), using the following protocol: 3 mL of methanol was added for conditioning followed by 2 mL of 0.1% trifluoroacetic acid (TFA) in H2O. The samples were then loaded onto each column followed by 4mL of 95:5: H2O:acetonitrile (ACN), 0.1% TFA. Samples were eluted with 1mL 80:20 ACN:H2O, 0.1% TFA. The samples were completely dried using a SpeedVac vacuum concentrator.

#### TMT-10 labeling of peptides

The dried tryptic peptides were dissolved with 500 mM HEPES (pH 8.5) and then labeled with 10-plex Tandem Mass Tag™ (TMT) reagents (Thermo Fisher Scientific) in 100% ACN. A ratio of TMT to peptide amount of 10:1 (w/w) was used (i.e., 500 μg of peptides labeled by 5 mg of TMT reagent). After incubation for one hour at room temperature, the reaction was terminated by adding 5% hydroxylamine for 15 minutes at room temperature. The TMT-labeled peptides were then acidified with 0.5% FA. Peptides labeled by different TMT reagents were then mixed, dried using a SpeedVac vacuum concentrator, reconstituted with 3% ACN, 0.1% FA and desalted again with C18 SPE.

#### Peptide fractionation and enrichment

The peptides were further fractionated using a reversed-phase Waters XBridge C18 column (250 mm × 4.6 mm column packed with 3.5-μm particles) on an Agilent 1200 HPLC System (solvent A: 5 mM ammonium formate, pH 10, 2% ACN; solvent B: 5 mM ammonium formate, pH 10, 90% ACN) operating at a flow rate of 1 mL/min [Anal. Chem. 2019, 91, 9, 5794–5801]. Peptides were separated by a gradient mixture from 0% B to 16% B in six minutes, 40% B in 60 minutes, 44% B in 4 min and ramped to 60% B in five minutes. The 60% B mixture was kept for 14 min. Fractions were collected into a 96 well plate during the fractionation run with a total of 96 fractions at the 1-minute time interval. The 96 fractions were subsequently concatenated into 24 fractions by combining 4 fractions that are 24 fractions apart (i.e., combining fractions #1, #25, #49, and #73; #2, #26, #50, and #74; and so on). For proteome analysis, 5% of each concatenated fraction was dried down and re-suspended in 2% acetonitrile, 0.1% formic acid to a peptide concentration of 0.1 mg/mL for LC-MS/MS analysis. The rest of the fractions (95%) were further concatenated into 12 fractions (i.e., by combining fractions #1 and #13; #3 and #15; and so on), dried down, and phosphopeptides enriched using immobilized metal affinity chromatography (IMAC).

#### Phosphopeptide enrichment using IMAC

The procedure for IMAC phosphopeptide enrichment has previously been reported here (Mertins et al., 2018). Briefly, Fe^3+^-NTA-agarose beads were freshly prepared using the Ni-NTA Superflow agarose beads (QIAGEN, #30410) for phosphopeptide enrichment. For each of the 12 fractions, peptides were reconstituted in 500 μL IMAC binding/wash buffer (80% ACN, 0.1% TFA) and incubated with 20 μL of the 50% bead suspension for 30 minutes at RT. After incubation, the beads were sequentially washed with 50 μL of the wash buffer (1X), 50 μL of 50% ACN, 0.1% TFA (1X), 50 μL of the wash buffer (1X), and 50 μL of 1% FA (1X) on the stage tip packed with 2 discs of Empore C18 material (Empore Octadecyl C18, 47 mm; Supleco, 66883-U). Phosphopeptides were eluted from the beads onto the C18 disc using 70 μL of the elution buffer (500 mM K2HPO4, pH 7.0). Sixty microliters of 50% ACN, 0.1% FA was used for the elution of phosphopeptides from the C18 stage tips after two washes with 100 μL of 1% FA. Samples were dried using a Speed-Vac and later reconstituted with 12 μL of 3% ACN, 0.1% FA for LC-MS/MS analysis.

#### LC-MS/MS Analysis

Lyophilized global and phosphorylated peptides were reconstituted in 12 μL of 0.1% FA with 2% ACN and 5 μL of the resulting sample was analyzed by LC-MS/MS using a Q-Exactive HF Quadrupole-Orbitrap Mass Spectrometer (Thermo Scientific) connected to a nanoACQUITY UPLC system (Waters Corp., Milford, MA) (buffer A: 0.1% FA with 3% ACN and buffer B: 0.1% FA in 90% ACN) as previously described (Tsai et al., 2020). Peptides were separated by a gradient mixture with an analytical column (75 μm i.d. × 25 cm) packed using 1.9- μm ReproSil C18 and with a column heater set at 50 °C. The analytical column was equilibrated to 98% buffer A and 2% buffer B and maintained at a constant column flow of 200 nL/min. Data were acquired in a data dependent mode with a full MS scan (350-1800 m/z) at a resolution of 60K with AGC setting set to 4×10^5^. The isolation window (quadrupole) for MS/MS was set at 0.7 m/z and optimal HCD fragmentation was performed at a normalized collision energy of 30% with AGC set as 1×10^5^ and a maximum ion injection time of 105 ms. The MS/MS spectra were acquired at a resolution of 50K. The dynamic exclusion time was set at 45 s.

#### MS Data Analysis

The raw MS/MS data were processed with MaxQuant (Cox and Mann, 2008; Tyanova et al., 2016a). The MS/MS spectra were searched against a human UniProt database (fasta file dated April 12, 2017 with 20,198 sequences). The search type was set to “Reporter ion MS2” for isobaric label measurements. A peptide search was performed with full tryptic digestion (Trypsin) and allowed a maximum of two missed cleavages. Carbamidomethyl (C) was set as a fixed modification; acetylation (protein N-term) and oxidation (M) were set as variable modifications for global proteome analysis. Acetylation (protein N-term), oxidation (M) and Phospho (STY) were set as variable modifications for phosphoproteome analysis. The false discovery rate (FDR) was set to 1% at the level of proteins, peptides, and modifications; no additional filtering was performed. The intensities of all ten TMT reporter ions were extracted from MaxQuant outputs and the abundances of TMT were firstly log2 transformed. The phosphoproteome data were further processed by the Ascore algorithm (Beausoleil et al., 2006) for phosphorylation site localization, and the top-scoring sequences were reported. The Perseus (Tyanova et al., 2016b) was used for statistical analyses.

#### Flow cytometry and fluorescence-activated cell sorting (FACS)

Flow cytometry was utilized to assess wild-type CXCR3 and CXCR3 mutant receptor cell surface expression in HEK293 cells. HEK293 cells seeded in six-well plates were transfected with wild-type CXCR3 or the indicated CXCR3 mutant using the calcium phosphate method. Forty-eight hours later, the cells were collected, washed with ice cold phosphate buffered saline (PBS), and subsequently centrifuged at 600 g for 4 minutes at 4 **°**C. Supernatant was aspirated and cells were resuspended in ice cold PBS and counted. 1E6 cells were transferred to a new tube and resuspended in 100 μL of blocking buffer (PBS + 3% FBS + 10mM EDTA + 5% Normal Human Serum) on ice for 5 to 10 minutes. PE conjugated anti-Human CD183 (CXCR3) antibody (R&D Systems, Minneapolis, MN) was added per the manufacturers guidelines and cells were incubated for 20 to 30 minutes at room temperature in the dark. Cells were centrifuged once more, supernatant aspirated, and fixed in 300 µL of 0.4% paraformaldehyde and were assessed using a BD LSRII flow cytometer. Flow cytometry was performed in the Duke Human Vaccine Institute Research Flow Cytometry Facility (Durham, NC). FACS was utilized to select Jurkat cells expressing wild-type CXCR3 or the indicated CXCR3 mutant. Following lentiviral transduction and subsequent puromycin selection, Jurkat cells were collected and washed in Hank’s Balanced Salt Solution (HBSS) (Gibco) with 2.5% FBS and 1.5 µM EDTA. Cells were then labelled with APC conjugated anti-Human CD183 (CXCR3) antibody (Biolegend, San Diego, CA) for 25 minutes on ice in the dark. Cells were then washed with HBSS with 2.5% FBS and 1.5 µM EDTA and resuspended with DNase. Cells were then strained through a sterile 30 µm filter and sorted on an Astrios (Beckman Coulter) sorter. Analyses were conducted with FlowJo version 10 software.

#### TGF-α shedding assay

G protein activity of various CXCR3 phosphorylation deficient mutants was assessed by the TGF-α shedding assay as previously described (Inoue et al., 2012). HEK293 cells were transiently transfected using Lipofectamine 2000 (Thermo Fisher Scientific) with wild-type CXCR3 or the indicated CXCR3 mutant receptor, modified TGF-α–containing alkaline phosphatase (AP-TGF-α), and the Gαi1 or Gαi3 subunit or the negative control GαΔc. 24 hours later, cells were detached and reseeded in HBSS with 5 mM HEPES in a clear-bottomed, white-walled, Costar 96-well plate (Corning Inc., Corning, NY). One hour later, cells were stimulated with the indicated concentration of CXCL11 for one hour. Conditioned medium (CM) containing the shed AP-TGF-α was transferred to a new 96-well plate. Both the cells and CM were treated with para-nitrophenylphosphate (p-NPP, 100 mM; Sigma-Aldrich, St. Louis, MO) substrate for one hour. The conversion of p-NPP to para-nitrophenol (p- NP) was measured at an optical density at 405 nm (OD405) in a BioTek Synergy Neo2 plate reader plate reader immediately after p-NPP addition and then after a 1-hour incubation. Gα activity was calculated by determining p-NP amounts by absorbance using the following equation:

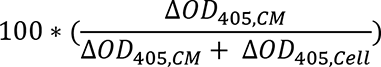

where ΔOD405 = OD405 at 1hr – OD405 at 0 hours and ΔOD405, cell and ΔOD405, CM represent the changes in absorbance after one hour in the cell and CM plates, respectively. Data were normalized to the negative control GαΔc.

#### Split luciferase and BRET assays

HEK293 cells seeded in six-well plates (∼750000 cells/well) were transfected with the appropriate constructs using the calcium-phosphate protocol. TRUPATH assays to assess G protein dissociation utilized wild-type CXCR3 or the indicated CXCR3 mutant, Gαi1-RLuc8, Gγ9-GFP2, and Gβ3 at equal amounts (Olsen *et al*., 2020). β-arrestin-2 recruitment was assessed using wild-type CXCR3 or the indicated CXCR3 mutant tagged with a C-terminal RLuc2 and a β-arrestin-2-mKO. Receptor internalization was assessed using wild-type CXCR3 or the indicated CXCR3 mutant tagged with a C-terminal RLuc2 and either a Myrpalm tagged mVenus to assess proximity to the cellular membrane, or a 2x-Fyve tagged mVenus to assess proximity to the early endosome. GRK recruitment was assessed using a split luciferase assay where wild-type CXCR3 or the indicated CXCR3 mutant was tagged with a SmBiT and GRK2, GRK3, GRK5, or GRK6 was tagged with a LgBiT.

Twenty-four hours after transfection, cells were washed with PBS, collected with trypsin, and plated onto clear-bottomed, white-walled, Costar 96-well plates at 50000 to 100000 cells/well in BRET medium (clear minimum essential medium (Gibco) supplemented with 2% fetal bovine serum, 10 mM HEPES, 1x GlutaMax (Gibco), and 1x Antibiotic-Antimycotic (Gibco)). The following day, media was removed, and cells were incubated at 37°C with 80 μL of HBSS supplemented with 20 mM HEPES and 3μM coelenterazine-400a (Cayman Chemical, Ann Arbor, MI) for TRUPATH or 3 µM coelenterazine h for all other BRET or split luciferase assays (Cayman Chemical, Ann Arbor, MI) for 10 to 15 minutes. For TRUPATH, plates were read with a BioTek Synergy Neo2 plate reader set at 37°C with a standard 400 nm emission filter and 510 nm long pass filter. For all other BRET assays, a standard 480 nm RLuc emission filter and 530 nm (for experiment using mVenus) or custom 542 nm (for experiments using mKO) long pass filter was utilized (Chroma Technology Co., Bellows Falls, VT). Cells were stimulated with either vehicle control (HBSS with 20 mM HEPES) or the indicated concentration of chemokine. All readings were performed using a kinetic protocol. For split luciferase experiments, plates were read before and after ligand treatment to calculate a change in luminescence after ligand stimulation and subsequently normalized to vehicle treatment. For BRET experiments, the BRET ratio was calculated by dividing the acceptor signal by the luciferase signal, and a net BRET ratio was calculated by normalizing to vehicle treatment.

#### Intramolecular Fluorescent Arsenical Hairpin (FlAsH) BRET of β-arrestin-2

FlAsH BRET experiments were carried out using a modified protocol as previously described (Lee *et al*., 2016; Strungs et al., 2019). FlAsH 3 serves as a negative control as insertion of the CCPGCC motif at this location significantly impacts β-arrestin recruitment to the receptor and does not demonstrate significant changes in BRET efficiency following ligand stimulation. HEK293 cells seeded in six-well plates were transfected with wild-type CXCR3 or the indicated CXCR3 mutant and FlAsH 1, 2, 3, 4, 5, or 6 using the calcium-phosphate protocol. Twenty-four hours after transfection, cells were washed with PBS, collected with trypsin, and plated onto clear-bottomed, rat-tail collagen coated, white-walled, Costar 96-well plates at 50000 to 100000 cells/well in supplemented MEM. The following day, cells were washed with 50 µL of HBSS and incubated in biarsenical labelling reagent FlAsH-EDT2 at a final concentration of 2.5 µM for 45 minutes at room temperature in the dark. Cells were then washed once with a 250 µM BAL wash buffer (2,3-dimercaptopropanol) and incubated with HBSS with 20 mM HEPES. Cells were stimulated by either vehicle control (HBSS with 20 mM HEPES) or chemokine for eight minutes. Immediately before reading the plate, cells were treated with coelenterazine h and read on a BioTek Synergy Neo2 plate reader set at 37°C using standard 480 nm and 530 nm emission filters. Net BRET values were calculated as described by averaging six consecutive BRET values and normalizing to vehicle control. Two-way ANOVA was performed at each FlAsH construct to determine if there was a significant ligand, receptor, or interaction term. If a significant interaction term was detected, Tukey’s post hoc testing was performed for multiple comparisons between receptor:ligand combinations at the specified FlAsH construct.

#### Molecular Dynamics

The model of the CXCR3 C-tail/ β-arrestin 2 complex was based on the structure of β-arrestin 1 in complex with the V2R C-tail (Shukla *et al*., 2013). The sequence of β-arrestin 2 was modified to match the isoform used in the FlAsH *in vitro* experiments [P29067]. The complexes were solvated (TIP3P water) and neutralized using a 0.15 M concentration of NaCl ions. Parameters for simulations were obtained from the Charmm36M forcefield (Huang et al., 2017). Simulations were run using the ACEMD3 engine (Harvey et al., 2009). All systems underwent a 40ns equilibration in conditions of constant pressure (NPT ensemble, pressure maintained with Berendsen barostat, 1.01325 bar), using a timestep of 2fs. During this stage mobility restraints were applied to the backbone. This was followed with 3 x 1.5µs of simulation for each system in conditions of constant volume (NVT ensemble) using a timestep of 4fs. For every simulation we used a temperature of 310K, maintained using the Langevin thermostat. Hydrogen bonds were restrained using the RATTLE algorithm. Non-bonded interactions were cut- off at a distance of 9Ȧ, with a smooth switching function applied at 7.5Ȧ. The interdomain rotation angle of β- arrestin 2 was analyzed using a script kindly provided by Naomi Latoracca (Latorraca *et al*., 2018). The angle was measured by comparing the displacement of the C-domain relative to the N-domain between the inactive (PDB code: 1G4R) and active βarr1 crystal structures (PDB code: 4JQI). Each simulation frame was aligned to the reference structures using the Cα atoms of the β-strands present within the N-domain, while the same atoms present in the C-domain were used to calculate the rotation angle. For each of the variants of the C-tail, we have phosphorylated all Ser and Thr residues present within the sequence. To study correlation of the interdomain rotation angle, and the distance between the studied probes and Arg8 (RLuc anchor point), we have utilized simulations of the L344X system (which in our setup meant that a C-tail was not included at all). Simulation data are shared on the open online resource GPCRmd (Rodriguez-Espigares et al., 2020) with the ID 1485.

#### Immunoblotting

Experiments were conducted as previously described (Smith et al., 2018b). Briefly, HEK293 cells were transiently transfected via the calcium-phosphate method with either wild-type CXCR3 or the indicated CXCR3 mutant. 48 hours after transfection, the cells were serum starved in minimum essential medium with 1% penicillin/streptomycin, 0.05% bovine serum albumin, and 5 mM HEPES for at least four hours, stimulated to a final concentration with 100 nM chemokine or vehicle control for 5, 30 or 60 minutes, subsequently washed once with ice-cold PBS, lysed in ice-cold radioimmunoprecipitation assay buffer (150 mM NaCl, 1% Nonidet P-40, 0.5% sodium deoxycholate, 0.1% sodium dodecyl sulfate (SDS), 25 mM Tris pH 7.4) containing the phosphatase inhibitor PhosSTOP (Roche, Basel, Switzerland) and protease inhibitor cOmplete EDTA free (Sigma-Aldrich, St. Louis, MO). Samples were then rotated for approximately 45 minutes at 4 °C and cleared of insoluble debris by centrifugation at 17000 g at 4 °C for 15 minutes, after which the supernatant was collected. Protein was resolved on SDS-10% polyacrylamide gels, transferred to nitrocellulose membranes, and immunoblotted with the indicated primary antibody overnight at 4 °C. phospho-ERK (Cell Signaling Technology) and total ERK (Millipore) were used to assess ERK activation. Peroxidase–conjugated polyclonal donkey anti-rabbit immunoglobulin (IgG) or polyclonal sheep anti-mouse IgG were used as secondary antibodies. Immune complexes on nitrocellulose membrane were imaged by SuperSignal enhanced chemiluminescent substrate (Thermo Fisher) using a ChemiDoc MP Imaging System (Bio-Rad). For quantification, phospho-ERK signal was normalized to total ERK signal using ImageLab (Bio-Rad) within the same immunoblot.

#### Confocal microscopy

HEK293 cells plated in rat-tail-collagen-coated 35 mm glass bottomed dishes (MatTek Corporation, Ashland, MA) were transiently transfected using FuGENE 6 with either wild-type CXCR3-GFP or the indicated CXCR3- GFP mutant and β-arrestin-2-RFP. 48 hours after transfection, the cells were serum starved for one hour prior to treatment with the indicated chemokine at 100 nM for 45 minutes at 37°C. The samples were then washed once with HBSS and fixed in a 6% formaldehyde solution for 30 minutes in the dark at room temperature. Cells were then washed four times with PBS and subsequently imaged with a Zeiss CSU-X1 spinning disk confocal microscope using the corresponding lasers to excite GFP (480 nm) and RFP (561 nm). Confocal images were arranged and analyzed using ImageJ (NIH, Bethesda, MD).

#### Generation of stably expressing CXCR3 Jurkats and Jurkat Chemotaxis

CXCR3 knock out (CXCR3-KO) Jurkat cells were generated using CRISPR-Cas9. CXCR3 guide RNA was developed using GAGTGACCACCAAGTGCTAAATGACG and GATGAAGTCTGGGAGGGCGAAA and inserted into a Cas9 containing plasmid backbone (PX459). Jurkat cells were transfected using Lipofectamine 2000 with the designed PX459 plasmid and CXCR3-KO Jurkats were selected using Puromycin and single clones were selected via limited dilution. CXCR3-KO was confirmed via flow cytometry. Stably expressing CXCR3 Jurkats were generated using lentiviral transduction. The wild-type or mutant CXCR3 were cloned into a pLenti plasmid backbone consisting of the receptor underneath a CMV promoter. HEK293 cells were transfected using calcium-phosphate with the pLenti receptor containing plasmid, envelope vector (pMD2.G), and packaging vector (psPAX2). 16 hours post-transfection, the HEK293 cell media was changed. 64 hours post transfection, the viral containing media was harvested, and virus was concentrated using the Lenti-X concentrator (Takara Bio, Japan) and viral titer was determined using qPCR per the manufacturer guidelines (ABM, Canada). CXCR3-KO Jurkats were transduced with virus via centrifugation at 1000 g for 90 minutes at a multiplicity of infection of 80-100 in the presence of polybrene at 8µg/mL. Cells expressing CXCR3 were sorted via FACS to obtain cells that express receptor to a similar degree. Chemotaxis assays were run in a 96 well format using the 5 µm ChemoTx chemotaxis system (Neuro Probe, Gaithersburg, MD). 750000 Jurkats were serum starved for at least four hours and placed in the chemotaxis system and allowed to migrate towards vehicle control or chemokine. Chemotaxis was measured using a previously described MTT labeling assay where the number of migrated cells is quantified by the reduction of MTT (Shi et al., 1993). Following chemotaxis, cells were labelled with a 0.5 mg/mL solution of MTT for four hours at 37 °C, subsequently lysed in 2 mM hydrochloric acid in isopropanol, and absorbance was read at an optical density of 570 nm. Chemotactic index was determined by measuring the absorbance of cells treated with chemokine to those treated with vehicle and normalized to the cell type with maximum chemotactic response.

#### Chemokines

Recombinant Human CXCL9, CXCL10, and CXCL11 (PeproTech) were diluted according to the manufacturer’s specifications, and aliquots were stored at −80 °C until needed for use.

### QUANTIFICATION AND STATISTICAL ANALYSIS

#### Statistical analyses

Data were analyzed in Excel (Microsoft, Redmond, WA) and graphed in Prism 9.0 (GraphPad, San Diego, CA). Dose-response curves were fitted to a log agonist versus stimulus with three parameters (span, baseline, and EC50) with the minimum baseline corrected to zero. For comparing ligands or receptors in concentration-response assays, a two-way ANOVA of ligand and concentration was conducted. Unless otherwise noted, statistical tests were two-sided and Tukey’s post hoc testing was performed for multiple comparisons or Dunnet’s testing was performed when comparisons were made to a reference condition. Statistical significance was shown on figures typically for the Emax of dose-response curves. In some cases, when applicable, statistical significance was shown on figures for EC50. Unless otherwise state, post hoc comparisons were made between CXCR3-WT and the denoted phosphorylation deficient receptor. Further details of statistical analysis and replicates are included in the figure captions. Experiments were not randomized, and investigators were not blinded to treatment conditions. Critical plate-based experiments were independently replicated by at least two different investigators when feasible.

## KEY RESOURCES TABLE

| REAGENT OR RESOURCE | SOURCE | IDENTIFIER |
| --- | --- | --- |
| **Antibodies** |  |  |
| Donkey polyclonal anti-rabbit IgG peroxidase conjugated | Rockland | Cat#611-7302; RRID:AB_219747 |
| Sheep polyclonal anti-mouse IgG peroxidase conjugated | Rockland | Cat#610-603-002; RRID:AB_219694 |
| Mouse monoclonal anti-phospho-p44/42 MAPK 1/2 (ERK1/2) (Thr202/Tyr204) | Cell Signaling Technologies | Cat#9106; RRID:AB_331768 |
| Rabbit polyclonal anti-MAPK 1/2 (ERK1/2) | Millipore Sigma | Cat#06-182; RRID:AB_310068 |
| Mouse monoclonal anti-human CD183 (CXCR3) PE conjugated | R&D Systems | Cat#FAB160P; RRID:AB_2086755 |
| Mouse monoclonal anti-human CD183 (CXCR3) APC conjugated | BioLegend | Cat#353707; RRID:AB_10962949 |
| **Bacterial Strains** |  |  |
| XL10-Gold Ultracompetent E. Coli | Agilent | Cat#200315 |
| **Chemicals, peptides, and recombinant proteins** |  |  |
| Recombinant Human CXCL9 | Peprtech | Cat#300-26 |
| Recombinant Human CXCL10 | Peprtech | Cat#300-12 |
| Recombinant Human CXCL11 | Peprtech | Cat#300-46 |
| GlutaMax | Gibco | Cat#35050061 |
| Antibiotic-Antimycotic | Gibco | Cat#15240062 |
| Fugene 6 | Promega | Cat#E2691 |
| Lipofectamine 2000 | Invitrogen | Cat#11668019 |
| para-Nitrophenyl Phosphate | Sigma-Aldrich | Cat#4876 |
| 2,3-dimercapto-1-propanol | Sigma-Aldrich | Cat#64046 |
| FIAsH-EDT2 | Santa Cruz Biotechnology | Cat#sc-363644 |
| Coelenterazine h | Cayman Chemical | Cat#16894 |
| Coelenterazine h | NanoLight Technology | Cat#301 |
| Coelenterazine 400a | Cayman Chemical | Cat#16157 |
| PhosSTOP | Sigma-Aldrich | Cat#4906845001 |
| cOmplete Protease Inhibitor Cocktail | Sigma-Aldrich | Cat#11697498001 |
| SuperSignal West Pico PLUS Chemiluminescent Substrate | Thermo Fischer Scientific | Cat#34580 |
| Lenti-X Concentrator | Takara Bio | Cat#631232 |
| Polybrene | Sigma-Aldrich | Cat#TR-1003 |
| **Critical commercial assays** |  |  |
| qPCR Lentivirus Titration Kit | Applied Biological Materials (ABM) | Cat#LV900 |
| QuikChange Lightning Site-Directed Mutagenesis Kit | Agilent | Cat#210518 |
| ChemoTx 5 μm Chemotaxis System | Neuroprobe | Cat#116-5 |
| **Mass Spectrometry Resources** |  |  |
| Ni-NTA Superflow Agarose Beads | Qiagen | Cat#30410 |
| BCA Protein Assay Kit | ThermoFisher Scientific | Cat#A53225 |
| TMT-11 reagent kit | ThermoFisher Scientific | Cat#A34808 |
| Trypsin | Promega | Cat#V5117 |
| Empore Octadecyl C18, 47 mm | Supleco | Cat#66883-U |
| Waters tC18 SepPak | Waters | Cat#WAT054925 |
| **Deposited Data** |  |  |
| Mass Spectrometry Proteomics Data | JPOST | JPST001599 (Accession Key 6844) |
| Molecular Dynamics Simulations | GPCRmd | 1485 |
| **Experimental Models: Cell Lines** |  |  |
| Human: 293 | ATCC | Cat#CRL-1573; RRID:CVCL_0045 |
| Human: 293 | ATCC | Cat#CRL-3216; RRID:CVCL_0063 |
| Human: 293 β-arrestin 1/2 Knock Out | Asuka Inoue | (Alvarez-Curto *et al.*, 2016) |
| Human: 293 GRK 2, 3, 5, 6 Knock Out | Asuka Inoue | (Pandey *et al.*, 2021b) |
| Human: Jurkat, Clone E6-1 | ATCC | Cat#TIB-152; RRID:CVCL_0367 |
| **Recombinant DNA** |  |  |
| CXCR3 | Rajagopal Lab | N/A |
| CXCR3-S355A/S356A | This work | N/A |
| CXCR3-T360A/S361A/ | This work | N/A |
| CXCR3-T360A/S361A/S364A/S366A | Rajagopal Lab (Smith *et al.*, 2021) | N/A |
| CXCR3-L344X | Rajagopal Lab (Smith *et al.*, 2017) | N/A |
| Gai1-RLuc8 | Bryan Roth Lab (Olsen *et al.*, 2020) | N/A |
| Gγ9-GFP2 | Bryan Roth Lab (Olsen *et al.*, 2020) | N/A |
| Gβ3 | Bryan Roth Lab (Olsen *et al.*, 2020) | N/A |
| CXCR3-RLucII | This work | N/A |
| CXCR3- S355A/S356A -RLucII | This work | N/A |
| CXCR3-T360A/S361A/-RLucII | This work | N/A |
| CXCR3-T360A/S361A/S364A/S366A-RLucII | This work | N/A |
| CXCR3-L344X-RLucII | This work | N/A |
| βarr2-mKO | Rajagopal Lab (Smith *et al.*, 2021) | N/A |
| Myrpalm-mVenus | Rajagopal Lab (Smith *et al.*, 2017) | N/A |
| 2x-Fyve-mvenus | Rajagopal Lab (Smith *et al.*, 2017) | N/A |
| CXCR3-GFP | Rajagopal Lab (Smith *et al.*, 2017) | N/A |
| CXCR3- S355A/S356A -GFP | This work | N/A |
| CXCR3-T360A/S361A/-GFP | This work | N/A |
| CXCR3-T360A/S361A/S364A/S366A-GFP | This work | N/A |
| CXCR3-L344X-GFP | This work | N/A |
| βarr2-RFP | Marc Caron Lab | N/A |
| βarr2-FIAsh biosensors 1-6 | Louis Luttrell Lab (Lee *et al.*, 2016) | N/A |
| CXCR3-SmBiT | This work | N/A |
| CXCR3- S355A/S356A -SmBiT | This work | N/A |
| CXCR3-T360A/S361A/-SmBiT | This work | N/A |
| CXCR3-T360A/S361A/S364A/S366A-SmBiT | This work | N/A |
| CXCR3-L344X-SmBiT | This work | N/A |
| GRK2-LgBiT | Asuka Inoue Lab | N/A |
| GRK3-LgBiT | Asuka Inoue Lab | N/A |
| GRK5-LgBiT | Asuka Inoue Lab | N/A |
| GRK6-LgBiT | Asuka Inoue Lab | N/A |
| pLenti-CXCR3 | This work | N/A |
| pLenti-CXCR3-S355A/S356A | This work | N/A |
| pLenti-CXCR3-T360A/S361A | This work | N/A |
| pLenti-CXCR3-T360A/S361A/S364A/S366A | This work | N/A |
| pLenti-CXCR3- L344X | This work | N/A |
| pMD2.G | Addgene | Cat#12259; RRID:Addgene_12259 |
| psPAX2 | Addgene | Cat#12260; RRID:Addgene_12260 |
| PX459 | Addgene | Cat#62988; RRID:Addgene_62988 |
| **Software and algorithms** |  |  |
| GraphPad Prism | GraphPad Software | https://www.graphpad.com/scientific-software/prism/ |
| ImageJ | (Schneider et al., 2012) | https://imagej.nih.gov/ij/ |
| Adobe Illustrator | Adobe | https://www.adobe.com/ |
| Excel | Microsoft | https://www.microsoft.com/en-us/microsoft-365/excel |
| Database for Annotation, Visualization, and Integrated Discovery (DAVID) | (Huang da et al., 2009a; b) | https://david.ncifcrf.gov/home.jsp |
| Kinase Enrichment Analysis | (Lachmann and Ma'ayan, 2009) | https://www.maayanlab.net/KEA2/ |
| Modification Motifs | (Bailey et al., 2006; Cheng et al., 2019) | https://meme-suite.org/meme/tools/momo |
| GProX | (Rigbolt et al., 2011) | http://gprox.sourceforge.net/ |
| MaxQuant | (Tyanova et al., 2016a) | https://www.maxquant.org/ |
| FlowJo | Becton, Dickinson & Company | https://www.flowjo.com/ |
| ImageLab | Bio-Rad | https://www.bio-rad.com/en-us/product/image-lab-software |
| BioRender | BioRender | https://biorender.com/ |
| GPCRdb | (Kooistra et al., 2021) | https://gpcrdb.org/ |

## SUPPLEMENTAL FIGURE TITLES AND LEGENDS

Supplemental Figure 1: Quantitation of CXCR3 C-terminal phosphopeptides and G protein activation, β- arrestin-2 recruitment, and surface expression of various CXCR3 phosphodeficient mutants. Related to Figure 1.

Abundance of singly phosphorylated (**A**) RDpSSWSETSEASYSGL, (**B**) RDSpSWSETSEASYSGL, and (**C**) RDSSWSEpTSEASYSGL peptide following stimulation with vehicle control or 100 nM of chemokine for 5 minutes. Mean ± SEM, n=2 technical replicates of 6 pooled biological replicates. P<.05, by one-way ANOVA, Tukey’s post hoc analysis. (**D-E**) Agonist dose-dependent TGF-α shedding assay of CXCR3-WT and various phosphorylation deficient mutants to assess Gαi1 and Gαi3 protein activation. Mean ± SEM, n=3 (**F-G**) β-arrestin- 2 recruitment of receptors treated with CXCL11. Mean ± SEM, n=3. (**H**) Surface expression of HEK293 cells transiently transfected with pcDNA 3.1 empty vector, CXCR3-WT, or denoted receptor as measured by flow cytometry. Mean ± SEM, n=3-5. *P<.05 by one-way ANOVA, Dunnett’s post hoc testing denotes comparisons to CXCR3-WT.

Supplemental Figure 2: G protein dissociation in β-arrestin-1/2 knockout cells and phosphomimetic mutants, β-arrestin-2 recruitment grouped by receptor. Related to Figure 2.

(**A-C**) G protein dissociation of receptors treated with chemokine in β-arrestin-1/2 knockout cells. (**D**) G protein dissociation of CXCR3-S355A/S356A in β-arrestin-1/2 knockout cells. (**E-G**) G protein dissociation of CXCR3- WT, CXCR3-S355A/S356A, or phosphomimetic mutant CXCR3-S355D/S356D treated with chemokine. (**H-L**) β- arrestin-2 recruitment of receptors treated with chemokine as grouped by receptor. For (**A-G**) TRUPATH and (**H-L**) β-arrestin-2 recruitment assays, data shown are the mean ± SEM, n=3-4. * denotes statistically significant differences between EMax of specified receptor and CXCR3-WT unless otherwise noted. Agonist dose-dependent data presented are the average of BRET values 5 to 10 minutes following ligand stimulation.

Supplemental Figure 3: Single color confocal microscopy, and orthogonal BRET based receptor internalization assay. Related to Figure 2.

(**A-E**) Confocal microscopy images of HEK293 cells transfected with Receptor-GFP and β-arrestin-2-RFP following treatment with vehicle control or 100 nM of the listed chemokine for 45 minutes. Images are representative of three biological replicates. (**F**) Schematic of BRET assay to detect receptor internalization away from the plasma membrane. (**G**) BRET data of receptor internalization using the acceptor Myrpalm-mVenus following stimulation with 100 nM of the listed chemokine in HEK293 cells. For internalization BRET (**G**) assays, data shown are the mean ± SEM, n=4. *P<.05 by two-way ANOVA, Dunnett’s post hoc testing between CXCR3- WT and all other receptor mutants at a specific ligand. Data presented are the average of BRET values from 20- 30 minutes following ligand stimulation.

Supplemental Figure 4: GRK5 and GRK6 Recruitment Data in wild-type HEK293 cells and GRK 2/3/5/6 knockout cells. Related to Figure 3.

Agonist dose-dependent data and kinetic data of maximum treatment dose of (**A-C**) GRK5 and (**G-I**) GRK6 recruitment as measured by a split nanoluciferase assay in HEK293 cells. Data are grouped by ligand treatment. Agonist dose-dependent (**D-F**) GRK5 recruitment and (**J-L**) GRK6 recruitment as measured in GRK 2/3/5/6 knock out cells. Mean ± SEM, n=3.

Supplemental Figure 5: GRK2 and GRK3 Recruitment Data as grouped by receptor and with phosphomimetic receptors. Related to Figure 3.

Agonist dose-dependent data and kinetic data of maximum treatment dose of (**A-E**) GRK2 recruitment and (**L-P**) GRK3 recruitment to listed receptor as measured by a split nanoluciferase assay. Data are grouped by receptor. (**F-K**) GRK2 and (**Q-V**) GRK3 recruitment to phosphomimetic receptors (**F-H**, **Q-S**) CXCR3- T360D/S361D, and (**I-K**, **T-V**) CXCR3-4xD. Mean ± SEM, n=3-4. * denotes statistically significant differences between EMax of specified receptor and CXCR3-WT. # denotes statistically significant differences between EC50 of specified receptor and CXCR3-WT.

Supplemental Figure 6: Source FlAsH Data. Related to Figure 3.

(**A-F**) FlAsH data as grouped by FlAsH probes 1-6 shown as heat maps. Intensity of color corresponds with change in net BRET. (**G**) Source data for all FlAsH construct, ligand, receptor, combinations. Mean ± SEM, n=5.

*P<.05 by two-way ANOVA for each FlAsH construct are shown in the heat maps to demonstrate statistical significance of a ligand effect, receptor effect, and or interaction. If a significant interaction term was identified, Tukey’s post hoc testing was performed, and comparisons are shown in panel (**G**).

**Supplemental** Figure 7**: Structural model of the construct used in the molecular dynamics simulations. Related to** Figure 4. This model highlights the location of FlAsH probes 1-5 (red spheres) on β-arrestin 2 and the N-terminal RLuc (highlighted in green). We demonstrate the transition between an inactivate state with a low interdomain rotation angle, and an activate state, with a high interdomain rotation angle.

Supplemental Figure 8: Approach and source data for mass spectrometry to assess the global phosphoproteome. Related to Figure 5.

(**A**) Schematic of experimental design of global phosphoproteomics experiments. (**B**) Venn diagram showing number of proteins and phosphoproteins identified. (**C**) Source data describing the proteome and phosphoproteome where Class I phosphorylation sites are defined as those with a localization probability of at least 0.75 (Olsen et al., 2006). (**D**) TMT labelling intensity across samples and pooled data demonstrating high degrees of replicability. (**E**) Heatmap and dendrogram of individual pooled technical replicates of HEK293 cells treated with vehicle control, CXCL9, CXCL10, or CXCL11 demonstrating technical replicates of specific treatments cluster together. (**F**) Visual representation of GProX clustering.

Supplemental Figure 9: Quantification of ERK1/2 phosphorylation at 5 minutes as grouped by receptor. Related to Figure 6.

(**A-E**) Quantification of western blots of phosphorylated ERK1/2 in HEK293 cells expressing the indicated receptor following stimulation with vehicle control or 100 nM of chemokine at five minutes. Mean ± SEM, n=4.

*P<.05 by two-way ANOVA. Tukey’s post hoc testing denotes comparisons between ligands. Chemokine treatments were significantly different from vehicle at all receptors.

Supplemental Figure 10: Jurkat chemotaxis as grouped by receptor and univariate analyses of chemotaxis data. Related to Figure 7.

(**A-F**) Normalized Jurkat chemotaxis data grouped by CXCR3-KO or receptor. Mean ± SEM, n=4. *P<.05 by two-way ANOVA, Tukey’s post hoc testing. (**G-I**) Univariate linear regression of G Protein activation, β-arrestin 2 recruitment, and MAPK activation versus chemotaxis. Shown are the best fit lines and 95% confidence intervals for each regression analysis. *P<.05 by F-test to determine if the slope of the best fit line is non-zero.

## REFERENCES

1. Allen, S.J., Crown, S.E., and Handel, T.M. (2007). Chemokine: receptor structure, interactions, and antagonism. Annu Rev Immunol 25, 787–820. 10.1146/annurev.immunol.24.021605.090529.

2. Alvarez-Curto, E., Inoue, A., Jenkins, L., Raihan, S.Z., Prihandoko, R., Tobin, A.B., and Milligan, G. (2016). Targeted Elimination of G Proteins and Arrestins Defines Their Specific Contributions to Both Intensity and Duration of G Protein-coupled Receptor Signaling. J Biol Chem 291, 27147–27159. 10.1074/jbc.M116.754887.

3. Baidya, M., Kumari, P., Dwivedi-Agnihotri, H., Pandey, S., Chaturvedi, M., Stepniewski, T.M., Kawakami, K., Cao, Y., Laporte, S.A., Selent, J., et al. (2020). Key phosphorylation sites in GPCRs orchestrate the contribution of beta-Arrestin 1 in ERK1/2 activation. EMBO Rep 21, e49886. 10.15252/embr.201949886.

4. Bailey, T.L., Williams, N., Misleh, C., and Li, W.W. (2006). MEME: discovering and analyzing DNA and protein sequence motifs. Nucleic Acids Res 34, W369–373. 10.1093/nar/gkl198.

5. Beausoleil, S.A., Villen, J., Gerber, S.A., Rush, J., and Gygi, S.P. (2006). A probability-based approach for high-throughput protein phosphorylation analysis and site localization. Nat Biotechnol 24, 1285–1292. 10.1038/nbt1240.

6. Bonacchi, A., Romagnani, P., Romanelli, R.G., Efsen, E., Annunziato, F., Lasagni, L., Francalanci, M., Serio, M., Laffi, G., Pinzani, M., et al. (2001). Signal transduction by the chemokine receptor CXCR3: activation of Ras/ERK, Src, and phosphatidylinositol 3-kinase/Akt controls cell migration and proliferation in human vascular pericytes. J Biol Chem 276, 9945–9954. 10.1074/jbc.M010303200.

7. Bouzo-Lorenzo, M., Santo-Zas, I., Lodeiro, M., Nogueiras, R., Casanueva, F.F., Castro, M., Pazos, Y., Tobin, A.B., Butcher, A.J., and Camina, J.P. (2016). Distinct phosphorylation sites on the ghrelin receptor, GHSR1a, establish a code that determines the functions of ss-arrestins. Sci Rep 6, 22495. 10.1038/srep22495.

8. Bradley, S.J., Molloy, C., Valuskova, P., Dwomoh, L., Scarpa, M., Rossi, M., Finlayson, L., Svensson, K.A., Chernet, E., Barth, V.N., et al. (2020). Biased M1-muscarinic-receptor-mutant mice inform the design of next-generation drugs. Nat Chem Biol 16, 240–249. 10.1038/s41589-019-0453-9.

9. Busillo, J.M., Armando, S., Sengupta, R., Meucci, O., Bouvier, M., and Benovic, J.L. (2010). Site-specific phosphorylation of CXCR4 is dynamically regulated by multiple kinases and results in differential modulation of CXCR4 signaling. J Biol Chem 285, 7805–7817. 10.1074/jbc.M109.091173.

10. Butcher, A.J., Prihandoko, R., Kong, K.C., McWilliams, P., Edwards, J.M., Bottrill, A., Mistry, S., and Tobin, A.B. (2011). Differential G-protein-coupled receptor phosphorylation provides evidence for a signaling bar code. J Biol Chem 286, 11506–11518. 10.1074/jbc.M110.154526.

11. Cahill, T.J., 3rd, Thomsen, A.R., Tarrasch, J.T., Plouffe, B., Nguyen, A.H., Yang, F., Huang, L.Y., Kahsai, A.W., Bassoni, D.L., Gavino, B.J., et al. (2017). Distinct conformations of GPCR-beta-arrestin complexes mediate desensitization, signaling, and endocytosis. Proc Natl Acad Sci U S A 114, 2562–2567. 10.1073/pnas.1701529114.

12. Chen, Q., Perry, N.A., Vishnivetskiy, S.A., Berndt, S., Gilbert, N.C., Zhuo, Y., Singh, P.K., Tholen, J., Ohi, M.D., Gurevich, E.V., et al. (2017). Structural basis of arrestin-3 activation and signaling. Nat Commun 8, 1427. 10.1038/s41467-017-01218-8.

13. Cheng, A., Grant, C.E., Noble, W.S., and Bailey, T.L. (2019). MoMo: discovery of statistically significant post-translational modification motifs. Bioinformatics 35, 2774–2782. 10.1093/bioinformatics/bty1058.

14. Chow, M.T., Ozga, A.J., Servis, R.L., Frederick, D.T., Lo, J.A., Fisher, D.E., Freeman, G.J., Boland, G.M., and Luster, A.D. (2019). Intratumoral Activity of the CXCR3 Chemokine System Is Required for the Efficacy of Anti-PD-1 Therapy. Immunity 50, 1498–1512 e1495. 10.1016/j.immuni.2019.04.010.

15. Colvin, R.A., Campanella, G.S., Manice, L.A., and Luster, A.D. (2006). CXCR3 requires tyrosine sulfation for ligand binding and a second extracellular loop arginine residue for ligand-induced chemotaxis. Mol Cell Biol 26, 5838–5849. 10.1128/MCB.00556-06.

16. Colvin, R.A., Campanella, G.S., Sun, J., and Luster, A.D. (2004). Intracellular domains of CXCR3 that mediate CXCL9, CXCL10, and CXCL11 function. J Biol Chem 279, 30219–30227. 10.1074/jbc.M403595200.

17. Corbisier, J., Gales, C., Huszagh, A., Parmentier, M., and Springael, J.Y. (2015). Biased signaling at chemokine receptors. J Biol Chem 290, 9542–9554. 10.1074/jbc.M114.596098.

18. Cox, J., and Mann, M. (2008). MaxQuant enables high peptide identification rates, individualized p.p.b.-range mass accuracies and proteome-wide protein quantification. Nat Biotechnol 26, 1367–1372. 10.1038/nbt.1511.

19. Dwivedi-Agnihotri, H., Chaturvedi, M., Baidya, M., Stepniewski, T.M., Pandey, S., Maharana, J., Srivastava, A., Caengprasath, N., Hanyaloglu, A.C., Selent, J., and Shukla, A.K. (2020). Distinct phosphorylation sites in a prototypical GPCR differently orchestrate beta-arrestin interaction, trafficking, and signaling. Sci Adv 6. 10.1126/sciadv.abb8368.

20. Eichel, K., Jullie, D., Barsi-Rhyne, B., Latorraca, N.R., Masureel, M., Sibarita, J.B., Dror, R.O., and von Zastrow, M. (2018). Catalytic activation of beta-arrestin by GPCRs. Nature 557, 381–386. 10.1038/s41586-018-0079-1.

21. Eiger, D.S., Boldizsar, N., Honeycutt, C.C., Gardner, J., and Rajagopal, S. (2021). Biased agonism at chemokine receptors. Cell Signal 78, 109862. 10.1016/j.cellsig.2020.109862.

22. Ferguson, S.S., Downey, W.E., 3rd, Colapietro, A.M., Barak, L.S., Menard, L., and Caron, M.G. (1996). Role of beta-arrestin in mediating agonist-promoted G protein-coupled receptor internalization. Science 271, 363–366. 10.1126/science.271.5247.363.

23. Ge, L., Ly, Y., Hollenberg, M., and DeFea, K. (2003). A beta-arrestin-dependent scaffold is associated with prolonged MAPK activation in pseudopodia during protease-activated receptor-2-induced chemotaxis. J Biol Chem 278, 34418–34426. 10.1074/jbc.M300573200.

24. Griffith, J.W., Sokol, C.L., and Luster, A.D. (2014). Chemokines and chemokine receptors: positioning cells for host defense and immunity. Annu Rev Immunol 32, 659–702. 10.1146/annurev-immunol-032713-120145.

25. Gurevich, V.V., and Gurevich, E.V. (2004). The molecular acrobatics of arrestin activation. Trends Pharmacol Sci 25, 105–111. 10.1016/j.tips.2003.12.008.

26. Gurevich, V.V., and Gurevich, E.V. (2019). GPCR Signaling Regulation: The Role of GRKs and Arrestins. Front Pharmacol 10, 125. 10.3389/fphar.2019.00125.

27. Harvey, M.J., Giupponi, G., and Fabritiis, G.D. (2009). ACEMD: Accelerating Biomolecular Dynamics in the Microsecond Time Scale. J Chem Theory Comput 5, 1632–1639. 10.1021/ct9000685.

28. Hauser, A.S., Attwood, M.M., Rask-Andersen, M., Schioth, H.B., and Gloriam, D.E. (2017). Trends in GPCR drug discovery: new agents, targets and indications. Nat Rev Drug Discov 16, 829–842. 10.1038/nrd.2017.178.

29. He, Q.T., Xiao, P., Huang, S.M., Jia, Y.L., Zhu, Z.L., Lin, J.Y., Yang, F., Tao, X.N., Zhao, R.J., Gao, F.Y., et al. (2021). Structural studies of phosphorylation-dependent interactions between the V2R receptor and arrestin-2. Nat Commun 12, 2396. 10.1038/s41467-021-22731-x.

30. Huang da, W., Sherman, B.T., and Lempicki, R.A. (2009a). Bioinformatics enrichment tools: paths toward the comprehensive functional analysis of large gene lists. Nucleic Acids Res 37, 1–13. 10.1093/nar/gkn923.

31. Huang da, W., Sherman, B.T., and Lempicki, R.A. (2009b). Systematic and integrative analysis of large gene lists using DAVID bioinformatics resources. Nat Protoc 4, 44–57. 10.1038/nprot.2008.211.

32. Huang, J., Rauscher, S., Nawrocki, G., Ran, T., Feig, M., de Groot, B.L., Grubmuller, H., and MacKerell, A.D., Jr. (2017). CHARMM36m: an improved force field for folded and intrinsically disordered proteins. Nat Methods 14, 71–73. 10.1038/nmeth.4067.

33. Inagaki, S., Ghirlando, R., Vishnivetskiy, S.A., Homan, K.T., White, J.F., Tesmer, J.J., Gurevich, V.V., and Grisshammer, R. (2015). G Protein-Coupled Receptor Kinase 2 (GRK2) and 5 (GRK5) Exhibit Selective Phosphorylation of the Neurotensin Receptor in Vitro. Biochemistry 54, 4320–4329. 10.1021/acs.biochem.5b00285.

34. Inoue, A., Ishiguro, J., Kitamura, H., Arima, N., Okutani, M., Shuto, A., Higashiyama, S., Ohwada, T., Arai, H., Makide, K., and Aoki, J. (2012). TGFalpha shedding assay: an accurate and versatile method for detecting GPCR activation. Nat Methods 9, 1021–1029. 10.1038/nmeth.2172.

35. Inoue, A., Raimondi, F., Kadji, F.M.N., Singh, G., Kishi, T., Uwamizu, A., Ono, Y., Shinjo, Y., Ishida, S., Arang, N., et al. (2019). Illuminating G-Protein-Coupling Selectivity of GPCRs. Cell 177, 1933–1947 e1925. 10.1016/j.cell.2019.04.044.

36. Jung, S.R., Kushmerick, C., Seo, J.B., Koh, D.S., and Hille, B. (2017). Muscarinic receptor regulates extracellular signal regulated kinase by two modes of arrestin binding. Proc Natl Acad Sci U S A 114, E5579–E5588. 10.1073/pnas.1700331114.

37. Kahsai, A.W., Pani, B., and Lefkowitz, R.J. (2018). GPCR signaling: conformational activation of arrestins. Cell Res 28, 783–784. 10.1038/s41422-018-0067-x.

38. Kaya, A.I., Perry, N.A., Gurevich, V.V., and Iverson, T.M. (2020). Phosphorylation barcode-dependent signal bias of the dopamine D1 receptor. Proc Natl Acad Sci U S A 117, 14139–14149. 10.1073/pnas.1918736117.

39. Keshava Prasad, T.S., Goel, R., Kandasamy, K., Keerthikumar, S., Kumar, S., Mathivanan, S., Telikicherla, D., Raju, R., Shafreen, B., Venugopal, A., et al. (2009). Human Protein Reference Database--2009 update. Nucleic Acids Res 37, D767–772. 10.1093/nar/gkn892.

40. Kim, Y.M., and Benovic, J.L. (2002). Differential roles of arrestin-2 interaction with clathrin and adaptor protein 2 in G protein-coupled receptor trafficking. J Biol Chem 277, 30760–30768. 10.1074/jbc.M204528200.

41. Kliewer, A., Schmiedel, F., Sianati, S., Bailey, A., Bateman, J.T., Levitt, E.S., Williams, J.T., Christie, M.J., and Schulz, S. (2019). Phosphorylation-deficient G-protein-biased mu-opioid receptors improve analgesia and diminish tolerance but worsen opioid side effects. Nat Commun 10, 367. 10.1038/s41467-018-08162-1.

42. Komolov, K.E., and Benovic, J.L. (2018). G protein-coupled receptor kinases: Past, present and future. Cell Signal 41, 17–24. 10.1016/j.cellsig.2017.07.004.

43. Kooistra, A.J., Mordalski, S., Pandy-Szekeres, G., Esguerra, M., Mamyrbekov, A., Munk, C., Keseru, G.M., and Gloriam, D.E. (2021). GPCRdb in 2021: integrating GPCR sequence, structure and function. Nucleic Acids Res 49, D335–D343. 10.1093/nar/gkaa1080.

44. Kufareva, I., Salanga, C.L., and Handel, T.M. (2015). Chemokine and chemokine receptor structure and interactions: implications for therapeutic strategies. Immunol Cell Biol 93, 372–383. 10.1038/icb.2015.15.

45. Kuo, P.T., Zeng, Z., Salim, N., Mattarollo, S., Wells, J.W., and Leggatt, G.R. (2018). The Role of CXCR3 and Its Chemokine Ligands in Skin Disease and Cancer. Front Med (Lausanne) 5, 271. 10.3389/fmed.2018.00271.

46. Lachmann, A., and Ma’ayan, A. (2009). KEA: kinase enrichment analysis. Bioinformatics 25, 684–686. 10.1093/bioinformatics/btp026.

47. Laporte, S.A., Oakley, R.H., Zhang, J., Holt, J.A., Ferguson, S.S., Caron, M.G., and Barak, L.S. (1999). The beta2-adrenergic receptor/betaarrestin complex recruits the clathrin adaptor AP-2 during endocytosis. Proc Natl Acad Sci U S A 96, 3712–3717. 10.1073/pnas.96.7.3712.

48. Latorraca, N.R., Masureel, M., Hollingsworth, S.A., Heydenreich, F.M., Suomivuori, C.M., Brinton, C., Townshend, R.J.L., Bouvier, M., Kobilka, B.K., and Dror, R.O. (2020). How GPCR Phosphorylation Patterns Orchestrate Arrestin-Mediated Signaling. Cell 183, 1813–1825 e1818. 10.1016/j.cell.2020.11.014.

49. Latorraca, N.R., Wang, J.K., Bauer, B., Townshend, R.J.L., Hollingsworth, S.A., Olivieri, J.E., Xu, H.E., Sommer, M.E., and Dror, R.O. (2018). Molecular mechanism of GPCR-mediated arrestin activation. Nature 557, 452–456. 10.1038/s41586-018-0077-3.

50. Lee, M.H., Appleton, K.M., Strungs, E.G., Kwon, J.Y., Morinelli, T.A., Peterson, Y.K., Laporte, S.A., and Luttrell, L.M. (2016). The conformational signature of beta-arrestin2 predicts its trafficking and signalling functions. Nature 531, 665–668. 10.1038/nature17154.

51. Lin, R., Choi, Y.H., Zidar, D.A., and Walker, J.K.L. (2018). beta-Arrestin-2-Dependent Signaling Promotes CCR4-mediated Chemotaxis of Murine T-Helper Type 2 Cells. Am J Respir Cell Mol Biol 58, 745–755. 10.1165/rcmb.2017-0240OC.

52. Luttrell, L.M., Wang, J., Plouffe, B., Smith, J.S., Yamani, L., Kaur, S., Jean-Charles, P.Y., Gauthier, C., Lee, M.H., Pani, B., et al. (2018). Manifold roles of beta-arrestins in GPCR signaling elucidated with siRNA and CRISPR/Cas9. Sci Signal 11. 10.1126/scisignal.aat7650.

53. Mann, A., Liebetrau, S., Klima, M., Dasgupta, P., Massotte, D., and Schulz, S. (2020). Agonist-induced phosphorylation bar code and differential post-activation signaling of the delta opioid receptor revealed by phosphosite-specific antibodies. Sci Rep 10, 8585. 10.1038/s41598-020-65589-7.

54. Mantovani, A. (1999). The chemokine system: redundancy for robust outputs. Immunol Today 20, 254–257. 10.1016/s0167-5699(99)01469-3.

55. Marsango, S., Ward, R.J., Jenkins, L., Butcher, A.J., Al Mahmud, Z., Dwomoh, L., Nagel, F., Schulz, S., Tikhonova, I.G., Tobin, A.B., and Milligan, G. (2022). Selective phosphorylation of threonine residues defines GPR84-arrestin interactions of biased ligands. J Biol Chem 298, 101932. 10.1016/j.jbc.2022.101932.

56. Mayer, D., Damberger, F.F., Samarasimhareddy, M., Feldmueller, M., Vuckovic, Z., Flock, T., Bauer, B., Mutt, E., Zosel, F., Allain, F.H.T., et al. (2019). Distinct G protein-coupled receptor phosphorylation motifs modulate arrestin affinity and activation and global conformation. Nat Commun 10, 1261. 10.1038/s41467-019-09204-y.

57. Mertins, P., Tang, L.C., Krug, K., Clark, D.J., Gritsenko, M.A., Chen, L., Clauser, K.R., Clauss, T.R., Shah, P., Gillette, M.A., et al. (2018). Reproducible workflow for multiplexed deep-scale proteome and phosphoproteome analysis of tumor tissues by liquid chromatography-mass spectrometry. Nat Protoc 13, 1632–1661. 10.1038/s41596-018-0006-9.

58. Miess, E., Gondin, A.B., Yousuf, A., Steinborn, R., Mosslein, N., Yang, Y., Goldner, M., Ruland, J.G., Bunemann, M., Krasel, C., et al. (2018). Multisite phosphorylation is required for sustained interaction with GRKs and arrestins during rapid mu-opioid receptor desensitization. Sci Signal 11. 10.1126/scisignal.aas9609.

59. Nobles, K.N., Xiao, K., Ahn, S., Shukla, A.K., Lam, C.M., Rajagopal, S., Strachan, R.T., Huang, T.Y., Bressler, E.A., Hara, M.R., et al. (2011). Distinct phosphorylation sites on the beta(2)-adrenergic receptor establish a barcode that encodes differential functions of beta-arrestin. Sci Signal 4, ra51. 10.1126/scisignal.2001707.

60. Nuber, S., Zabel, U., Lorenz, K., Nuber, A., Milligan, G., Tobin, A.B., Lohse, M.J., and Hoffmann, C. (2016). beta-Arrestin biosensors reveal a rapid, receptor-dependent activation/deactivation cycle. Nature 531, 661–664. 10.1038/nature17198.

61. Oakley, R.H., Laporte, S.A., Holt, J.A., Barak, L.S., and Caron, M.G. (1999). Association of beta-arrestin with G protein-coupled receptors during clathrin-mediated endocytosis dictates the profile of receptor resensitization. J Biol Chem 274, 32248–32257. 10.1074/jbc.274.45.32248.

62. Olsen, J.V., Blagoev, B., Gnad, F., Macek, B., Kumar, C., Mortensen, P., and Mann, M. (2006). Global, in vivo, and site-specific phosphorylation dynamics in signaling networks. Cell 127, 635–648. 10.1016/j.cell.2006.09.026.

63. Olsen, R.H.J., DiBerto, J.F., English, J.G., Glaudin, A.M., Krumm, B.E., Slocum, S.T., Che, T., Gavin, A.C., McCorvy, J.D., Roth, B.L., and Strachan, R.T. (2020). TRUPATH, an open-source biosensor platform for interrogating the GPCR transducerome. Nat Chem Biol 16, 841–849. 10.1038/s41589-020-0535-8.

64. Pack, T.F., Orlen, M.I., Ray, C., Peterson, S.M., and Caron, M.G. (2018). The dopamine D2 receptor can directly recruit and activate GRK2 without G protein activation. J Biol Chem 293, 6161–6171. 10.1074/jbc.RA117.001300.

65. Pandey, S., Kumari, P., Baidya, M., Kise, R., Cao, Y., Dwivedi-Agnihotri, H., Banerjee, R., Li, X.X., Cui, C.S., Lee, J.D., et al. (2021a). Intrinsic bias at non-canonical, beta-arrestin-coupled seven transmembrane receptors. Mol Cell 81, 4605–4621 e4611. 10.1016/j.molcel.2021.09.007.

66. Pandey, S., Kumari, P., Baidya, M., Kise, R., Cao, Y., Dwivedi-Agnihotri, H., Banerjee, R., Li, X.X., Cui, C.S., Lee, J.D., et al. (2021b). Intrinsic bias at non-canonical, beta-arrestin-coupled seven transmembrane receptors. Mol Cell. 10.1016/j.molcel.2021.09.007.

67. Pupo, A.S., Duarte, D.A., Lima, V., Teixeira, L.B., Parreiras, E.S.L.T., and Costa-Neto, C.M. (2016). Recent updates on GPCR biased agonism. Pharmacol Res 112, 49–57. 10.1016/j.phrs.2016.01.031.

68. Rajagopal, S., Bassoni, D.L., Campbell, J.J., Gerard, N.P., Gerard, C., and Wehrman, T.S. (2013). Biased agonism as a mechanism for differential signaling by chemokine receptors. The Journal of biological chemistry 288, 35039–35048. 10.1074/jbc.M113.479113.

69. Ribas, C., Penela, P., Murga, C., Salcedo, A., Garcia-Hoz, C., Jurado-Pueyo, M., Aymerich, I., and Mayor, F., Jr. (2007). The G protein-coupled receptor kinase (GRK) interactome: role of GRKs in GPCR regulation and signaling. Biochim Biophys Acta 1768, 913–922. 10.1016/j.bbamem.2006.09.019.

70. Rigbolt, K.T., Vanselow, J.T., and Blagoev, B. (2011). GProX, a user-friendly platform for bioinformatics analysis and visualization of quantitative proteomics data. Mol Cell Proteomics 10, O110 007450. 10.1074/mcp.O110.007450.

71. Rodriguez-Espigares, I., Torrens-Fontanals, M., Tiemann, J.K.S., Aranda-Garcia, D., Ramirez-Anguita, J.M., Stepniewski, T.M., Worp, N., Varela-Rial, A., Morales-Pastor, A., Medel-Lacruz, B., et al. (2020). GPCRmd uncovers the dynamics of the 3D-GPCRome. Nat Methods 17, 777–787. 10.1038/s41592-020-0884-y.

72. Sato, P.Y., Chuprun, J.K., Schwartz, M., and Koch, W.J. (2015). The evolving impact of g protein-coupled receptor kinases in cardiac health and disease. Physiol Rev 95, 377–404. 10.1152/physrev.00015.2014.

73. Scarpa, M., Molloy, C., Jenkins, L., Strellis, B., Budgett, R.F., Hesse, S., Dwomoh, L., Marsango, S., Tejeda, G.S., Rossi, M., et al. (2021). Biased M1 muscarinic receptor mutant mice show accelerated progression of prion neurodegenerative disease. Proc Natl Acad Sci U S A 118. 10.1073/pnas.2107389118.

74. Schneider, C.A., Rasband, W.S., and Eliceiri, K.W. (2012). NIH Image to ImageJ: 25 years of image analysis. Nat Methods 9, 671–675. 10.1038/nmeth.2089.

75. Sente, A., Peer, R., Srivastava, A., Baidya, M., Lesk, A.M., Balaji, S., Shukla, A.K., Babu, M.M., and Flock, T. (2018). Molecular mechanism of modulating arrestin conformation by GPCR phosphorylation. Nat Struct Mol Biol 25, 538–545. 10.1038/s41594-018-0071-3.

76. Shahabuddin, S., Ji, R., Wang, P., Brailoiu, E., Dun, N., Yang, Y., Aksoy, M.O., and Kelsen, S.G. (2006). CXCR3 chemokine receptor-induced chemotaxis in human airway epithelial cells: role of p38 MAPK and PI3K signaling pathways. Am J Physiol Cell Physiol 291, C34–39. 10.1152/ajpcell.00441.2005.

77. Shi, Y., Kornovski, B.S., Savani, R., and Turley, E.A. (1993). A rapid, multiwell colorimetric assay for chemotaxis. J Immunol Methods 164, 149–154. 10.1016/0022-1759(93)90307-s.

78. Shukla, A.K., Manglik, A., Kruse, A.C., Xiao, K., Reis, R.I., Tseng, W.C., Staus, D.P., Hilger, D., Uysal, S., Huang, L.Y., et al. (2013). Structure of active beta-arrestin-1 bound to a G-protein-coupled receptor phosphopeptide. Nature 497, 137–141. 10.1038/nature12120.

79. Shukla, A.K., Violin, J.D., Whalen, E.J., Gesty-Palmer, D., Shenoy, S.K., and Lefkowitz, R.J. (2008). Distinct conformational changes in beta-arrestin report biased agonism at seven-transmembrane receptors. Proc Natl Acad Sci U S A 105, 9988–9993. 10.1073/pnas.0804246105.

80. Smith, J.S., Alagesan, P., Desai, N.K., Pack, T.F., Wu, J.H., Inoue, A., Freedman, N.J., and Rajagopal, S. (2017). C-X-C Motif Chemokine Receptor 3 Splice Variants Differentially Activate Beta-Arrestins to Regulate Downstream Signaling Pathways. Molecular pharmacology 92, 136–150. 10.1124/mol.117.108522.

81. Smith, J.S., Nicholson, L.T., Suwanpradid, J., Glenn, R.A., Knape, N.M., Alagesan, P., Gundry, J.N., Wehrman, T.S., Atwater, A.R., Gunn, M.D., et al. (2018a). Biased agonists of the chemokine receptor CXCR3 differentially control chemotaxis and inflammation. Science Signaling 11, eaaq1075. 10.1126/scisignal.aaq1075.

82. Smith, J.S., Nicholson, L.T., Suwanpradid, J., Glenn, R.A., Knape, N.M., Alagesan, P., Gundry, J.N., Wehrman, T.S., Atwater, A.R., Gunn, M.D., et al. (2018b). Biased agonists of the chemokine receptor CXCR3 differentially control chemotaxis and inflammation. Sci Signal 11. 10.1126/scisignal.aaq1075.

83. Smith, J.S., Pack, T.F., Inoue, A., Lee, C., Zheng, K., Choi, I., Eiger, D.S., Warman, A., Xiong, X., Ma, Z., et al. (2021). Noncanonical scaffolding of Galphai and beta-arrestin by G protein-coupled receptors. Science 371. 10.1126/science.aay1833.

84. Smith, J.S., and Rajagopal, S. (2016). The beta-Arrestins: Multifunctional Regulators of G Protein-coupled Receptors. J Biol Chem 291, 8969–8977. 10.1074/jbc.R115.713313.

85. Smith, J.S., Rajagopal, S., and Atwater, A.R. (2018c). Chemokine Signaling in Allergic Contact Dermatitis: Toward Targeted Therapies. Dermatitis 29, 179–186. 10.1097/DER.0000000000000391.

86. Stevers, L.M., Sijbesma, E., Botta, M., MacKintosh, C., Obsil, T., Landrieu, I., Cau, Y., Wilson, A.J., Karawajczyk, A., Eickhoff, J., et al. (2018). Modulators of 14-3-3 Protein-Protein Interactions. J Med Chem 61, 3755–3778. 10.1021/acs.jmedchem.7b00574.

87. Strungs, E.G., Luttrell, L.M., and Lee, M.H. (2019). Probing Arrestin Function Using Intramolecular FlAsH- BRET Biosensors. Methods Mol Biol 1957, 309–322. 10.1007/978-1-4939-9158-7_19.

88. Sun, Y., Cheng, Z., Ma, L., and Pei, G. (2002). Beta-arrestin2 is critically involved in CXCR4-mediated chemotaxis, and this is mediated by its enhancement of p38 MAPK activation. J Biol Chem 277, 49212–49219. 10.1074/jbc.M207294200.

89. Tobin, A.B. (2008). G-protein-coupled receptor phosphorylation: where, when and by whom. Br J Pharmacol 153 *Suppl 1*, S167–176. 10.1038/sj.bjp.0707662.

90. Tsai, C.F., Smith, J.S., Krajewski, K., Zhao, R., Moghieb, A.M., Nicora, C.D., Xiong, X., Moore, R.J., Liu, T., Smith, R.D., et al. (2019). Tandem Mass Tag Labeling Facilitates Reversed-Phase Liquid Chromatography-Mass Spectrometry Analysis of Hydrophilic Phosphopeptides. Anal Chem 91, 11606–11613. 10.1021/acs.analchem.9b01814.

91. Tsai, C.F., Zhao, R., Williams, S.M., Moore, R.J., Schultz, K., Chrisler, W.B., Pasa-Tolic, L., Rodland, K.D., Smith, R.D., Shi, T., et al. (2020). An Improved Boosting to Amplify Signal with Isobaric Labeling (iBASIL) Strategy for Precise Quantitative Single-cell Proteomics. Mol Cell Proteomics 19, 828–838. 10.1074/mcp.RA119.001857.

92. Tyanova, S., Temu, T., and Cox, J. (2016a). The MaxQuant computational platform for mass spectrometry-based shotgun proteomics. Nat Protoc 11, 2301–2319. 10.1038/nprot.2016.136.

93. Tyanova, S., Temu, T., Sinitcyn, P., Carlson, A., Hein, M.Y., Geiger, T., Mann, M., and Cox, J. (2016b). The Perseus computational platform for comprehensive analysis of (prote)omics data. Nat Methods 13, 731–740. 10.1038/nmeth.3901.

94. Watanabe, N., and Osada, H. (2012). Phosphorylation-dependent protein-protein interaction modules as potential molecular targets for cancer therapy. Curr Drug Targets 13, 1654–1658. 10.2174/138945012803530035.

95. Watanabe, Y., Yoshizawa, A.C., Ishihama, Y., and Okuda, S. (2021). The jPOST Repository as a Public Data Repository for Shotgun Proteomics. Methods Mol Biol 2259, 309–322. 10.1007/978-1-0716-1178-4_20.

96. Xiao, K., Shenoy, S.K., Nobles, K., and Lefkowitz, R.J. (2004). Activation-dependent conformational changes in {beta}-arrestin 2. J Biol Chem 279, 55744–55753. 10.1074/jbc.M409785200.

97. Yang, F., Yu, X., Liu, C., Qu, C.X., Gong, Z., Liu, H.D., Li, F.H., Wang, H.M., He, D.F., Yi, F., et al. (2015). Phospho-selective mechanisms of arrestin conformations and functions revealed by unnatural amino acid incorporation and (19)F-NMR. Nat Commun 6, 8202. 10.1038/ncomms9202.

98. Yang, Z., Yang, F., Zhang, D., Liu, Z., Lin, A., Liu, C., Xiao, P., Yu, X., and Sun, J.P. (2017). Phosphorylation of G Protein-Coupled Receptors: From the Barcode Hypothesis to the Flute Model. Mol Pharmacol 92, 201–210. 10.1124/mol.116.107839.

99. Zheng, K., Smith, J.S., Eiger, D.S., Warman, A., Choi, I., Honeycutt, C.C., Boldizsar, N., Gundry, J.N., Pack, T.F., Inoue, A., et al. (2022). Biased agonists of the chemokine receptor CXCR3 differentially signal through Galphai:beta-arrestin complexes. Sci Signal 15, eabg5203. 10.1126/scisignal.abg5203.

100. Zhou, X.E., He, Y., de Waal, P.W., Gao, X., Kang, Y., Van Eps, N., Yin, Y., Pal, K., Goswami, D., White, T.A., et al. (2017). Identification of Phosphorylation Codes for Arrestin Recruitment by G Protein-Coupled Receptors. Cell 170, 457–469 e413. 10.1016/j.cell.2017.07.002.

101. Zlotnik, A., and Yoshie, O. (2012). The chemokine superfamily revisited. Immunity 36, 705–716. 10.1016/j.immuni.2012.05.008.

