## Supplementary Figures for "Phosphorylation barcodes direct biased chemokine signaling at CXCR3"

### SUPPLEMENTAL FIGURE 1

Supplemental Figure 1

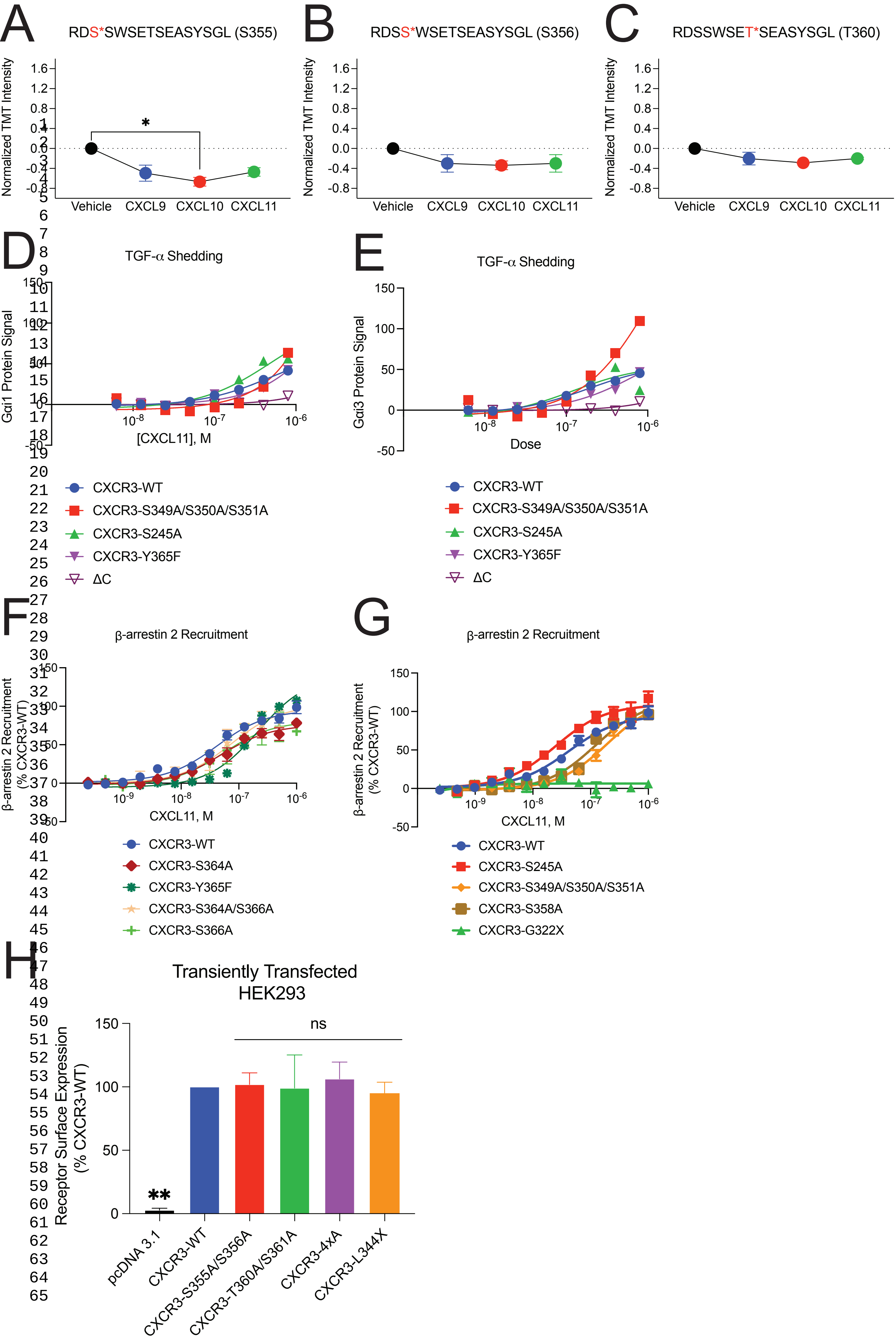

SUPPLEMENTAL FIGURE 2
Supplemental Figure 2

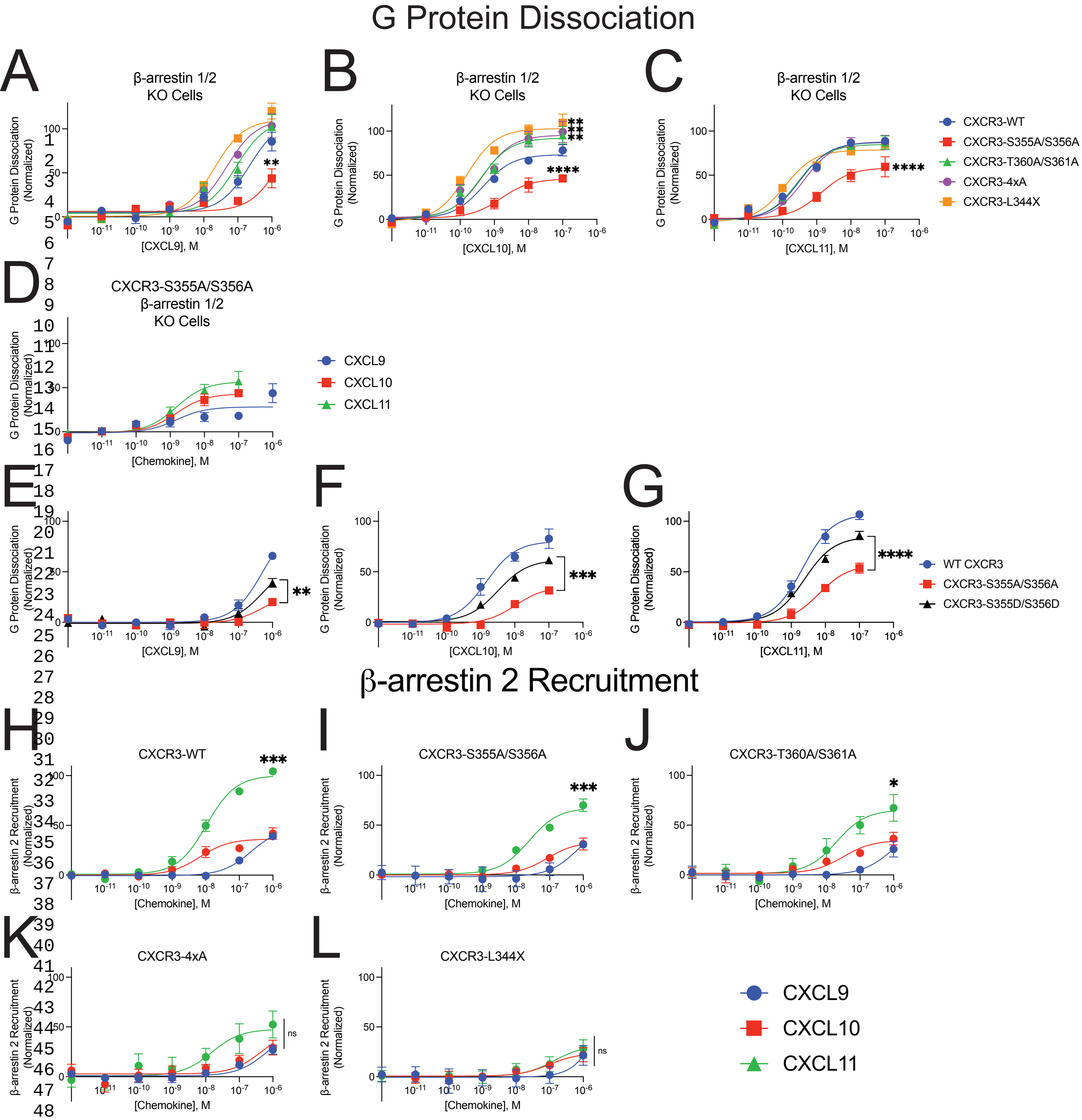

SUPPLEMENTAL FIGURE 3
Supplemental Figure 3

### Receptor Internalization

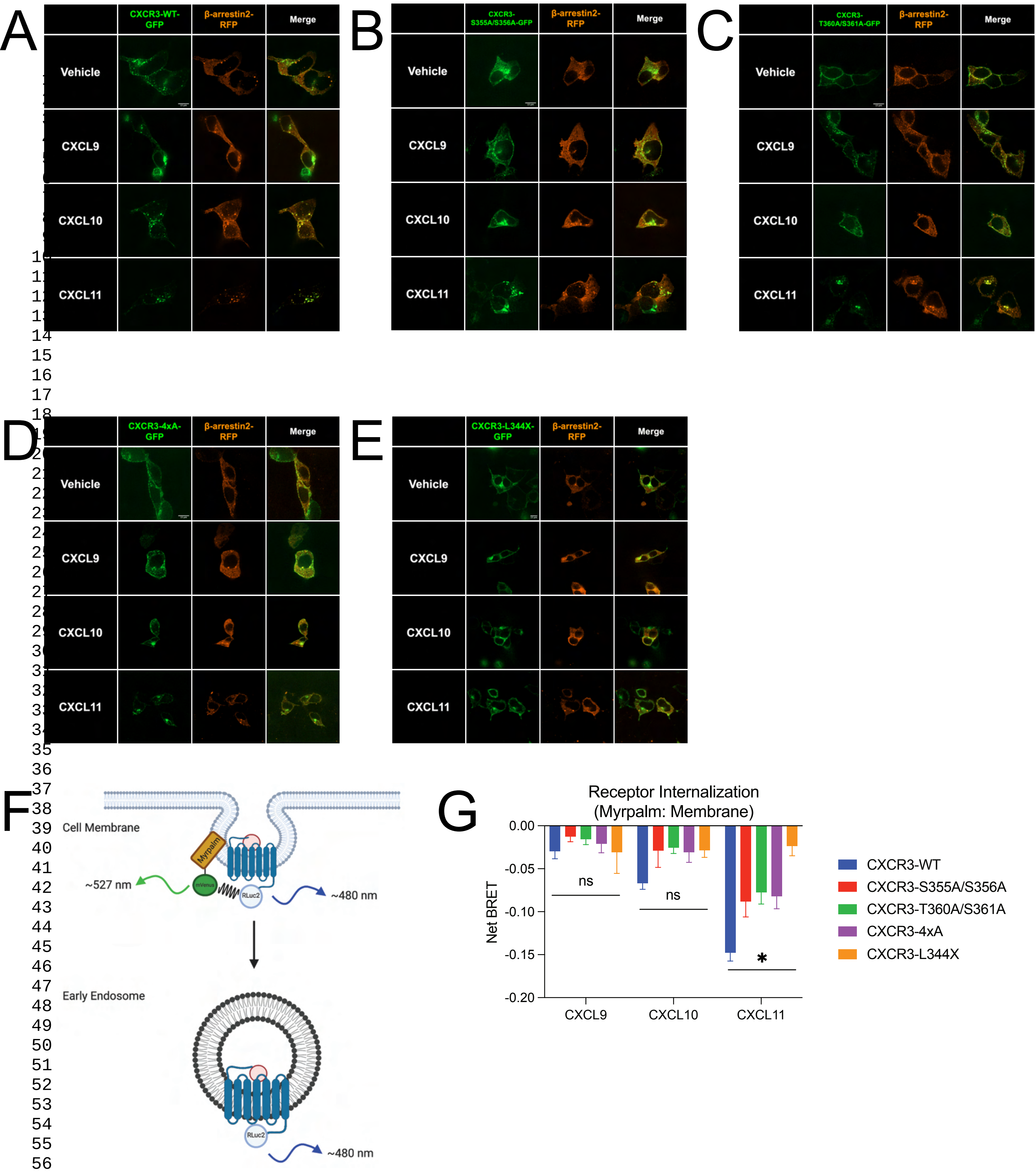

SUPPLEMENTAL FIGURE 4

Supplemental Figure 4

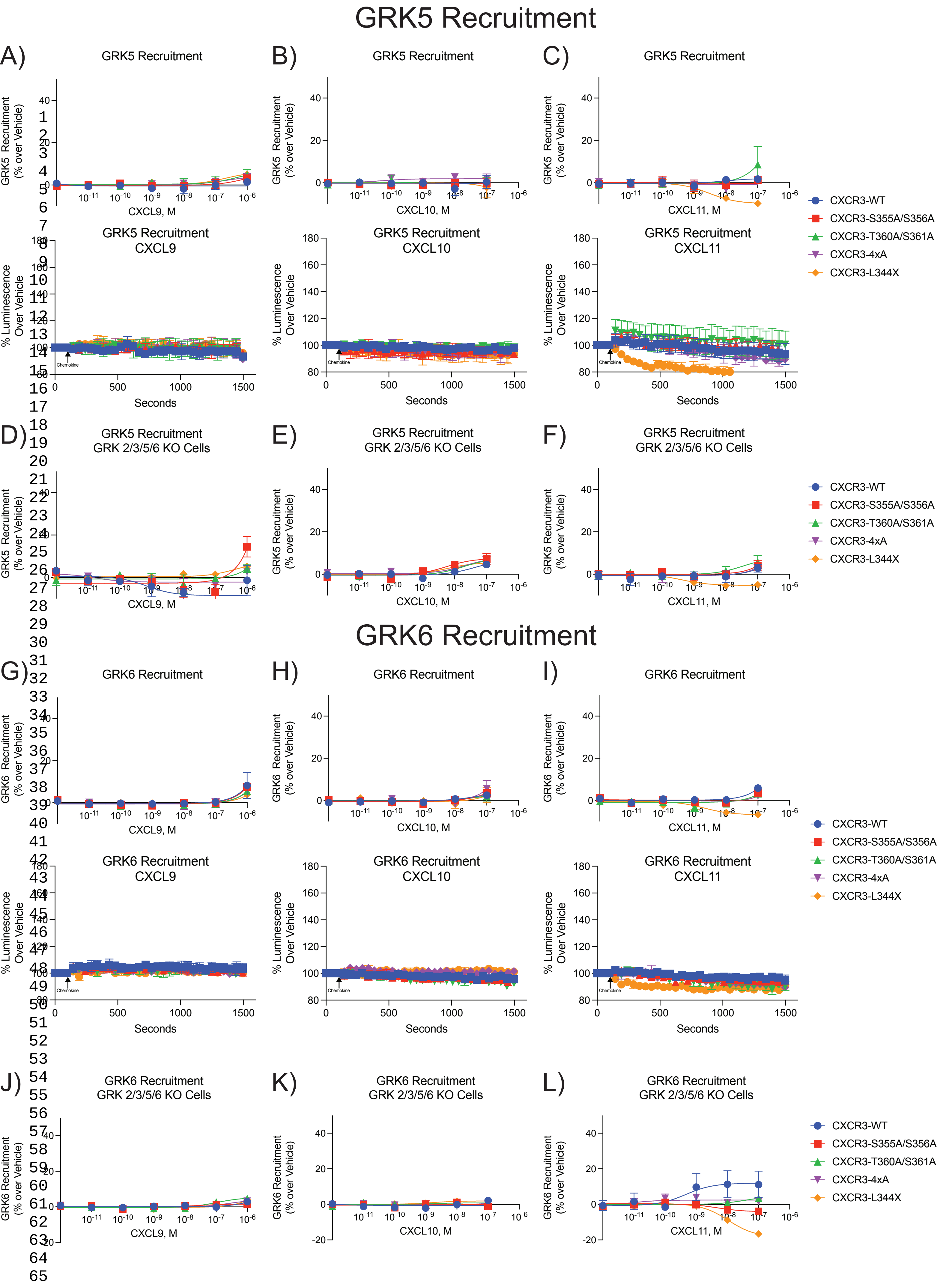

### SUPPLEMENTAL FIGURE 5

Supplemental Figure 5

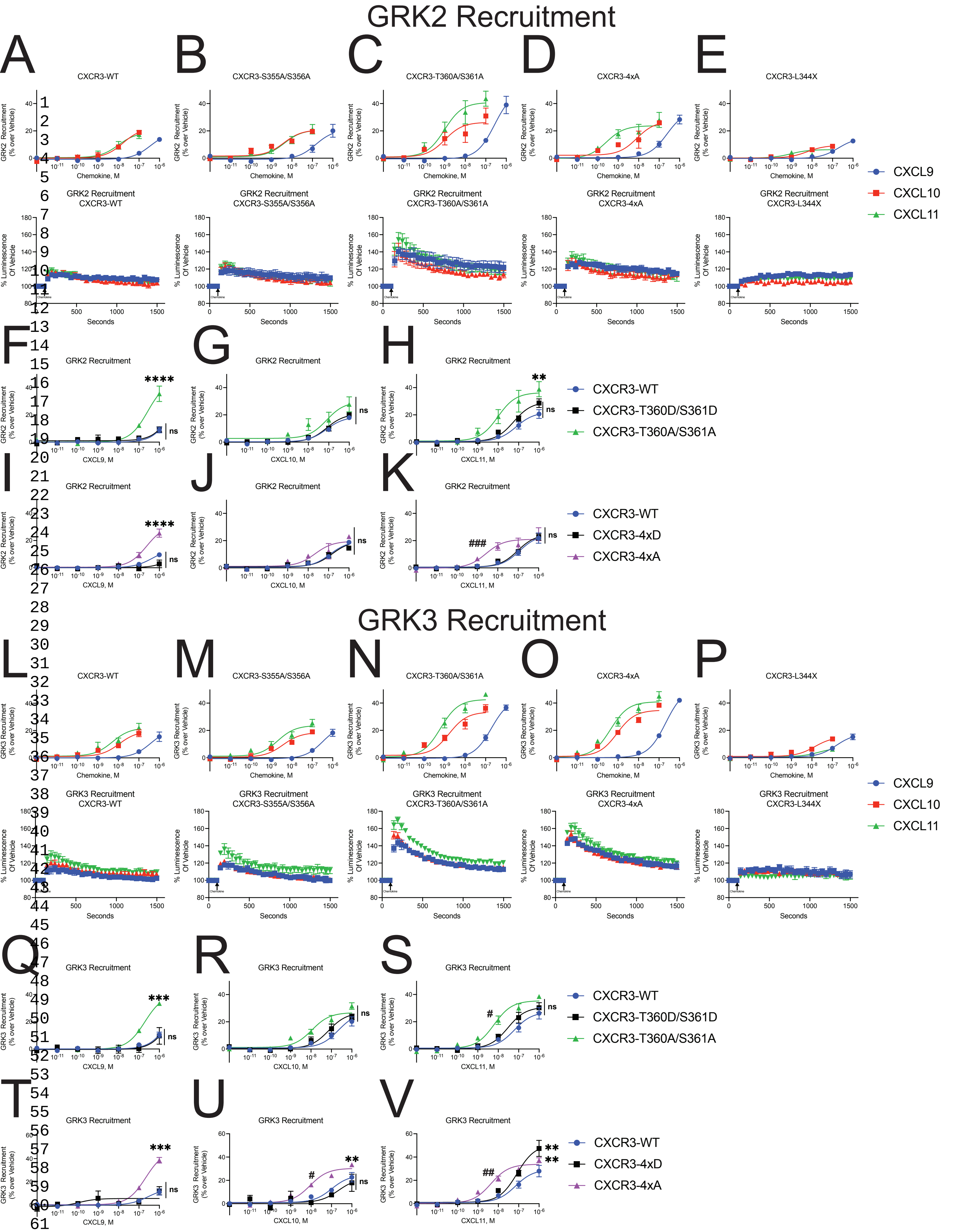

### SUPPLEMENTAL FIGURE 6

Supplemental Figure 6

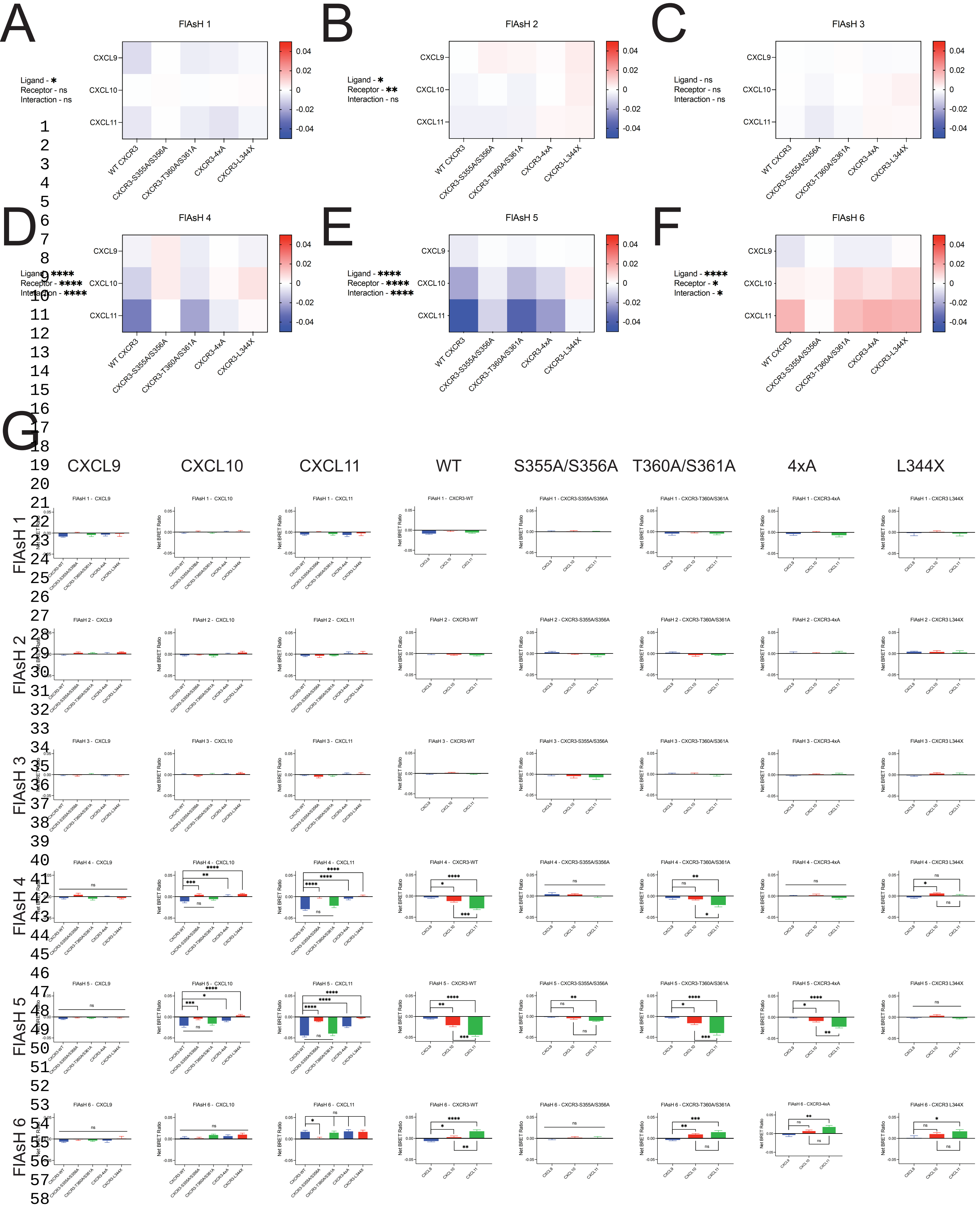

### SUPPLEMENTAL FIGURE 7

Supplemental Figure 7

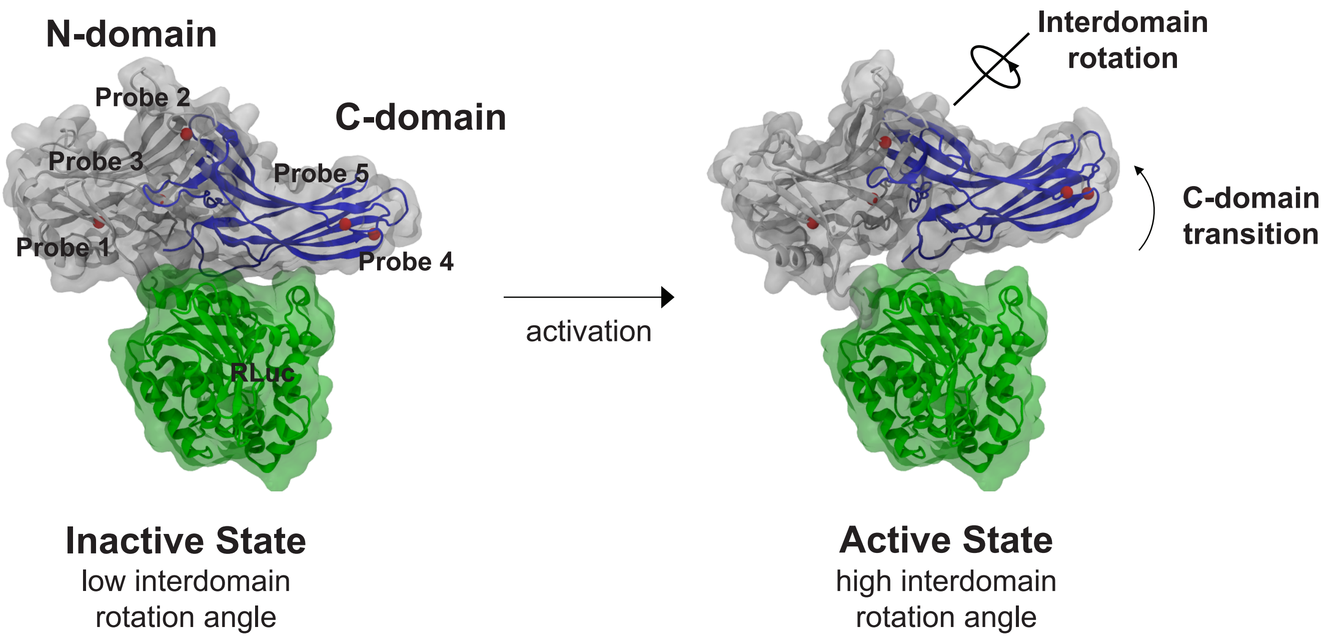

### SUPPLEMENTAL FIGURE 8

Supplemental Figure 8

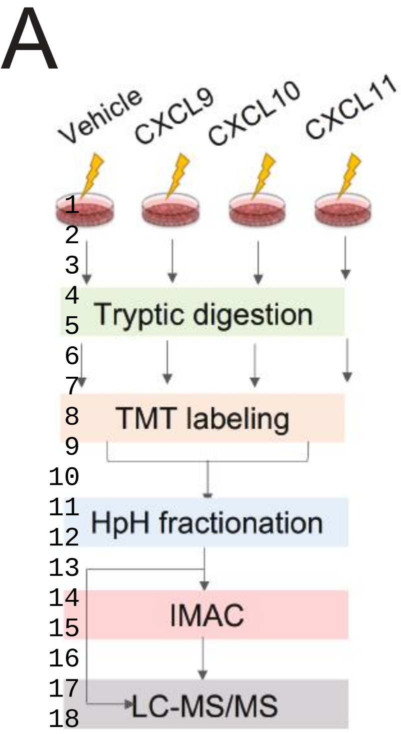


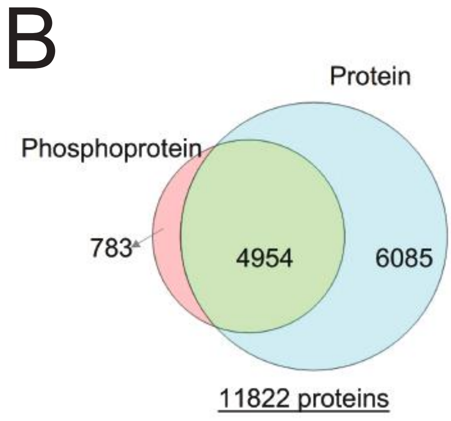


C

| Proteome | |
|---|---|
| Total Peptides | 159739 |
| Proteins | 11038 |

| Phosphoproteome | |
|---|---|
| Total Peptides | 29992 |
| Phosphopeptides | 26556 |
| Phosphorylation Sites | 22620 |
| Phosphoproteins | 5736 |
| Class 1 Phosphorylation Sites | 16448 |
| S | 18388 |
| T | 4112 |
| Y | 120 |

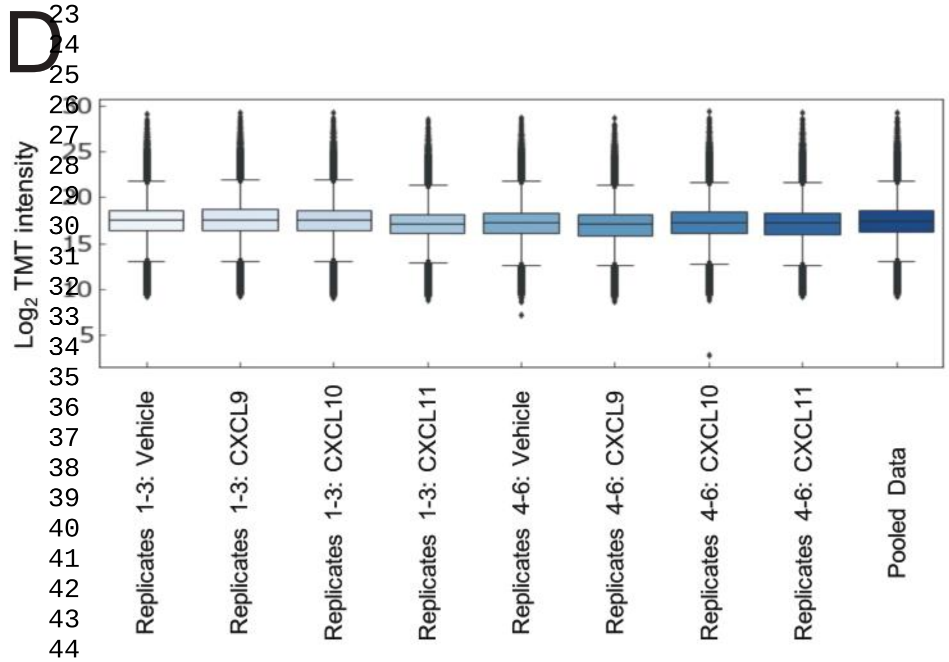


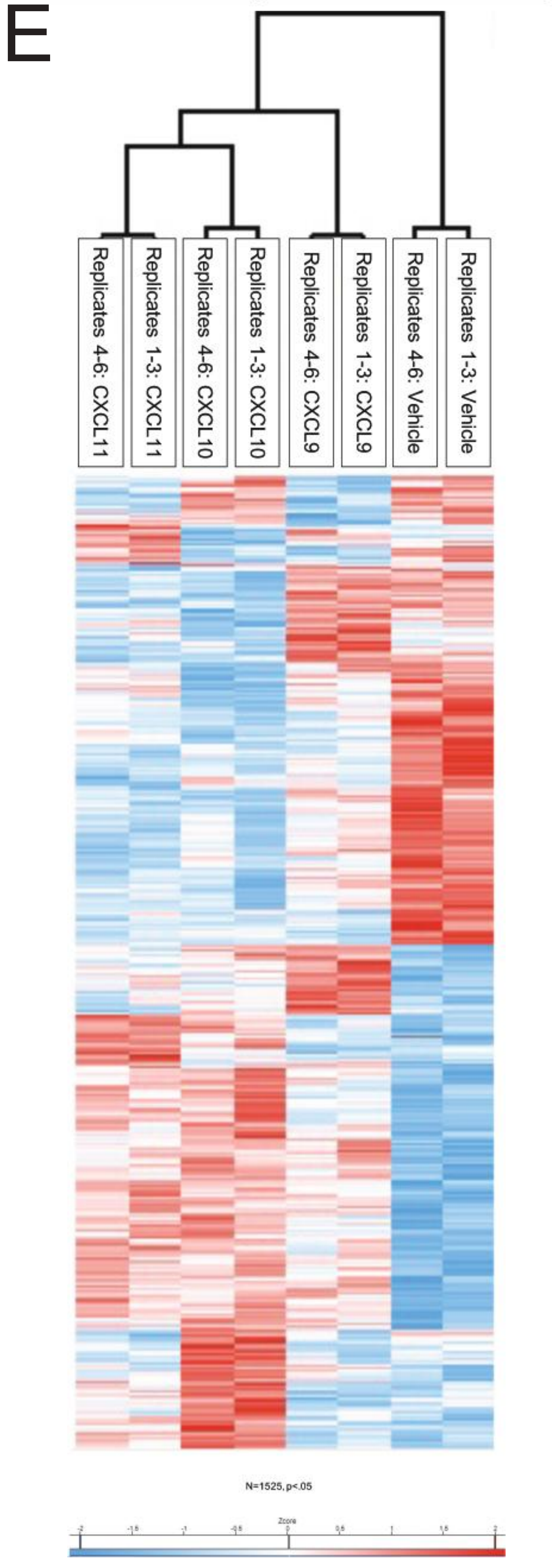


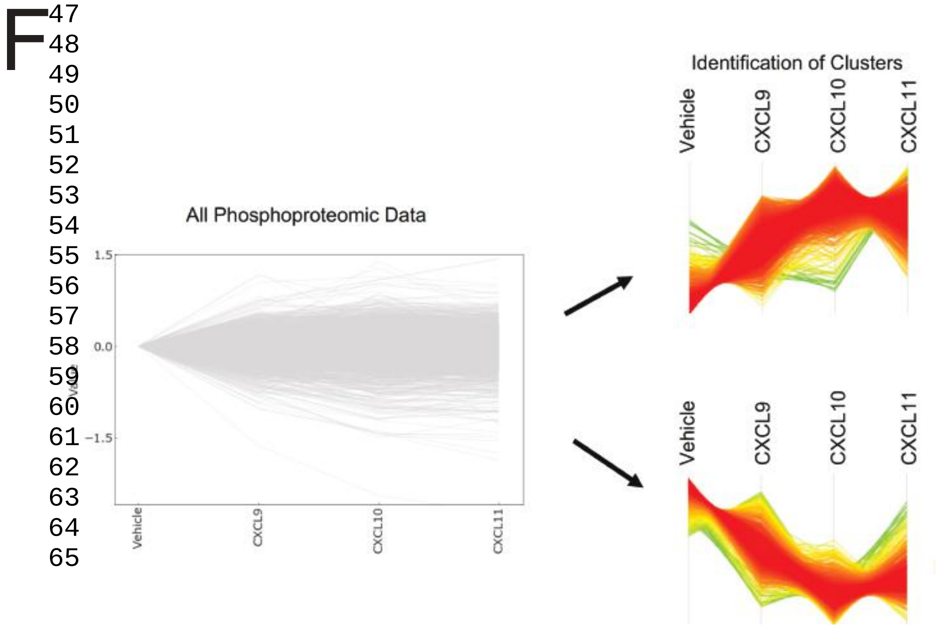

### SUPPLEMENTAL FIGURE 9

Supplemental Figure 9

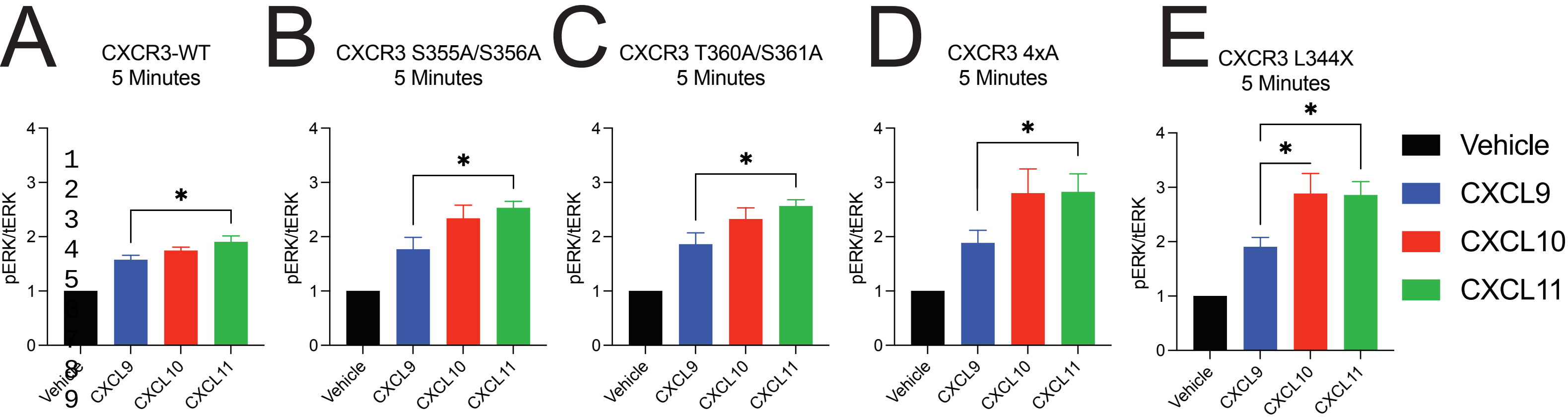

SUPPLEMENTAL FIGURE 10

Supplemental Figure 10

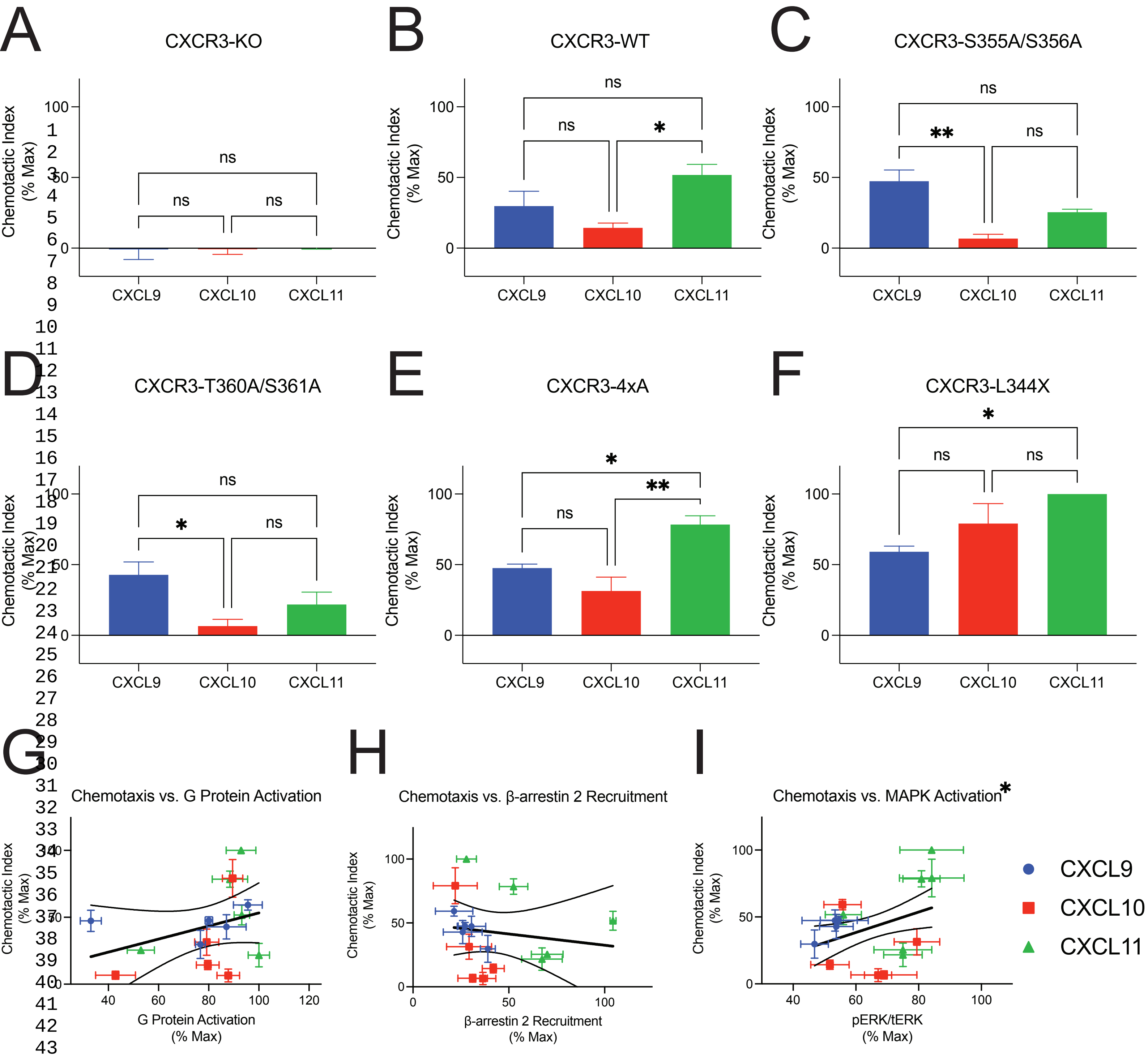
